# Mannose metabolism inhibition sensitizes acute myeloid leukemia cells to cytarabine and FLT3 inhibitor therapy by modulating fatty acid metabolism to drive ferroptotic cell death

**DOI:** 10.1101/2022.05.16.492042

**Authors:** Keith Woodley, Laura S Dillingh, George Giotopoulos, Pedro Madrigal, Kevin M Rattigan, Celine Philippe, Vilma Dembitz, Aoife M.S Magee, Ryan Asby, Louie N van de Lagemaat, Christopher Mapperley, Sophie C James, Jochen H.M Prehn, Konstantinos Tzelepis, Kevin Rouault-Pierre, George S Vassiliou, Kamil R Kranc, G Vignir Helgason, Brian J.P Huntly, Paolo Gallipoli

## Abstract

Resistance to standard and novel therapies remains the main obstacle to cure in acute myeloid leukemia (AML) and is often driven by metabolic adaptations which are therapeutically actionable. Here we identify inhibition of mannose-6-phosphate isomerase (MPI), the first enzyme in the mannose metabolism pathway, as a sensitizer to both cytarabine and FLT3 inhibitors across multiple AML models. Mechanistically, we identify a connection between mannose metabolism and fatty acid metabolism, that is mediated via preferential activation of the ATF6 arm of the unfolded protein response (UPR). This in turn leads to cellular accumulation of polyunsaturated fatty acids, lipid peroxidation and ferroptotic cell death in AML cells. Our findings provide further support to the role of rewired metabolism in AML therapy resistance, unveil a novel connection between two apparently independent metabolic pathways and support further efforts to achieve eradication of therapy-resistant AML cells by sensitizing them to ferroptotic cell death.

## Introduction

Resistance to therapy leading to disease relapse is the most frequent cause of treatment failure in acute myeloid leukemia (AML)^1^ and commonly results from the emergence of genetic mutations, often within the therapeutic target^2,3^. However clinical and preclinical studies, in both solid^4^ and hematological malignancies^5^ have shown that early non-genetic adaptations might allow some cancer cells to withstand therapeutic stress, while allowing the development of a fully resistant phenotype through either established adaptive changes or the subsequent acquisition of genetic mutations^6^. Adaptive changes can be driven by metabolic rewiring and metabolism has emerged as a therapeutically actionable vulnerability in AML^7^, where specific metabolic adaptations arising as a result of driver mutations or in response to therapy have been reported^8–13^.

D-Mannose is the 2-epimer of glucose and is a six-carbon sugar mostly used by cells for production of glycoconjugates rather than as an energy source. Mannose is transported into cells via transporters of the *SLC2A* group (GLUT) family and once within the cell is phosphorylated by hexokinase (HK) to produce mannose-6-phosphate (Man-6-P), which can mostly be either interconverted to the glycolytic intermediate fructose-6-phosphate (Fru-6-P) by mannose-6-phosphate isomerase (MPI) or directed into N-glycosylation via phosphomannomutase (PMM2)^14^. Since mannose plasma concentration is about 100-fold less than glucose concentration, the bidirectional MPI enzyme also plays a role in channelling glucose into the mannose metabolism (MM) pathway to feed the production of GDP-Mannose, a sugar donor for N-glycosylation reactions. Indeed, most of the mannose in N-glycans derives from glucose^15^ and as a result MPI sits at a branching point between N-glycosylation and energy metabolism pathways, such as glycolysis and the hexosamine biosynthetic pathway (HBP) (Extended Fig.1A). MPI plays an important role in embryonic development as *Mpi* knockout mice are embryonic lethal^16^ and only hypomorphs rather than completely inactivating mutations are described in patients^17,18^. However partial loss of MPI function causes a congenital disorder of glycosylation (CDG; MPI-CDG) in humans which is successfully treated with mannose supplementation^19^ while hypomorphic *Mpi* mice are viable^20^. Recent reports have highlighted a role for MPI in cancer cell proliferation or survival through varied mechanisms; regulation of TP53 O-linked-N-acetylglucosaminylation (O-GlcNAcylation) via modulation of HBP activity^21^, reduction of cell surface receptor glycosylation and signalling^22^ or modulating the susceptibility of several cancers to the toxic effects of high mannose diets^23^. Regardless of the mechanism, high MPI activity appears to provide a survival advantage in several cancer types. Interestingly, MPI was amongst the top drop-out genes in our published CRISPR-Cas9 screen aiming to identify sensitizers to the clinical grade FLT3-tyrosine kinase inhibitor (TKI) AC220 (quizartinib) in AML carrying activating FLT3 internal tandem duplication (ITD) mutations^9^ and more recently was identified amongst 81 United States Food and Drug Administration (FDA) druggable genetic dependencies of a murine model of FLT3^ITD^ AML^24^.

We therefore aimed to test if inhibition of MPI and MM sensitizes AML cells to FLT3-TKI and standard chemotherapy and characterise the mechanistic consequences of MPI inhibition.

## Results

### High MPI expression correlates with worse prognosis in AML and a gene signature associated with enhanced oxidative phosphorylation and fatty acid metabolism

Given its high millimolar plasma concentration glucose generates the majority of cellular Man-6-P via MPI ^15^. Our previous^9^ liquid chromatography-mass spectrometry (LC/MS) experiments with uniformly-labelled ^13^Carbon(U-^13^C_6_)-glucose in FLT3^ITD^ mutant cells treated with FLT3-TKI show that upon FLT3 inhibition, glucose metabolism is blocked at the level of Fru-6-P (Extended Data Fig.1A). This suggests that while glycolysis is inhibited, metabolic pathways branching from Fru-6-P, such as MM are still active and might play a cytoprotective role. Corroborating this notion, it is noticeable that other enzymes downstream of MPI in MM, i.e. GMPPB, were depleted in our published screen^9^ (Extended Data Fig.1A). Interestingly analysis of 2 published gene expression profiles of newly diagnosed AML cases associated with patient outcome shows that higher levels of *MPI* expression correlated with significant or borderline significant worse patient outcome (Fig.1A and Extended Data Fig.1B). *MPI* expression levels are higher in AML mononuclear cells (MNC) compared to normal bone marrow MNC (Fig.1B and Extended Data Fig.1C-D) and particularly in FLT3^ITD^ compared to FLT3^WT^ AML (Fig.1C and Extended Data Fig.1E) suggesting that MPI might have a prominent role in this AML subtype. When specifically focusing on FLT3^ITD^ mutant patients we observed a trend towards improved survival in patients expressing low *MPI* levels (Extended Data Fig.1F). However this was not significant possibly because of the small numbers. Gene-set enrichment analysis (GSEA) from several published gene expression datasets of AML samples at diagnosis highlighted oxidative phosphorylation, fatty acid oxidation and MYC targets gene signatures as being consistently upregulated in patients expressing high levels of *MPI* (Fig.1D and Extended Data Fig.1G). Enhanced oxidative phosphorylation and fatty acid oxidation are known features of AML cells resistant to both current and novel therapies^10,13^. Further supporting the role of MPI in therapy resistance, analysis of paired diagnosis and relapsed samples following standard chemotherapy from 2 independent datasets^25–27^ shows that *MPI* expression levels increase upon relapse (Fig.1E). Taken together these data suggest that MM and particularly *MPI* expression levels might play a role in resistance to both FLT3-TKI and standard therapies in AML. While this phenotype might be particularly associated with FLT3^IT^D mutations, higher *MPI* expression also correlates with gene signatures associated with drug resistance in AML beyond the presence of FLT3^IT^D mutations.

**Figure 1.**
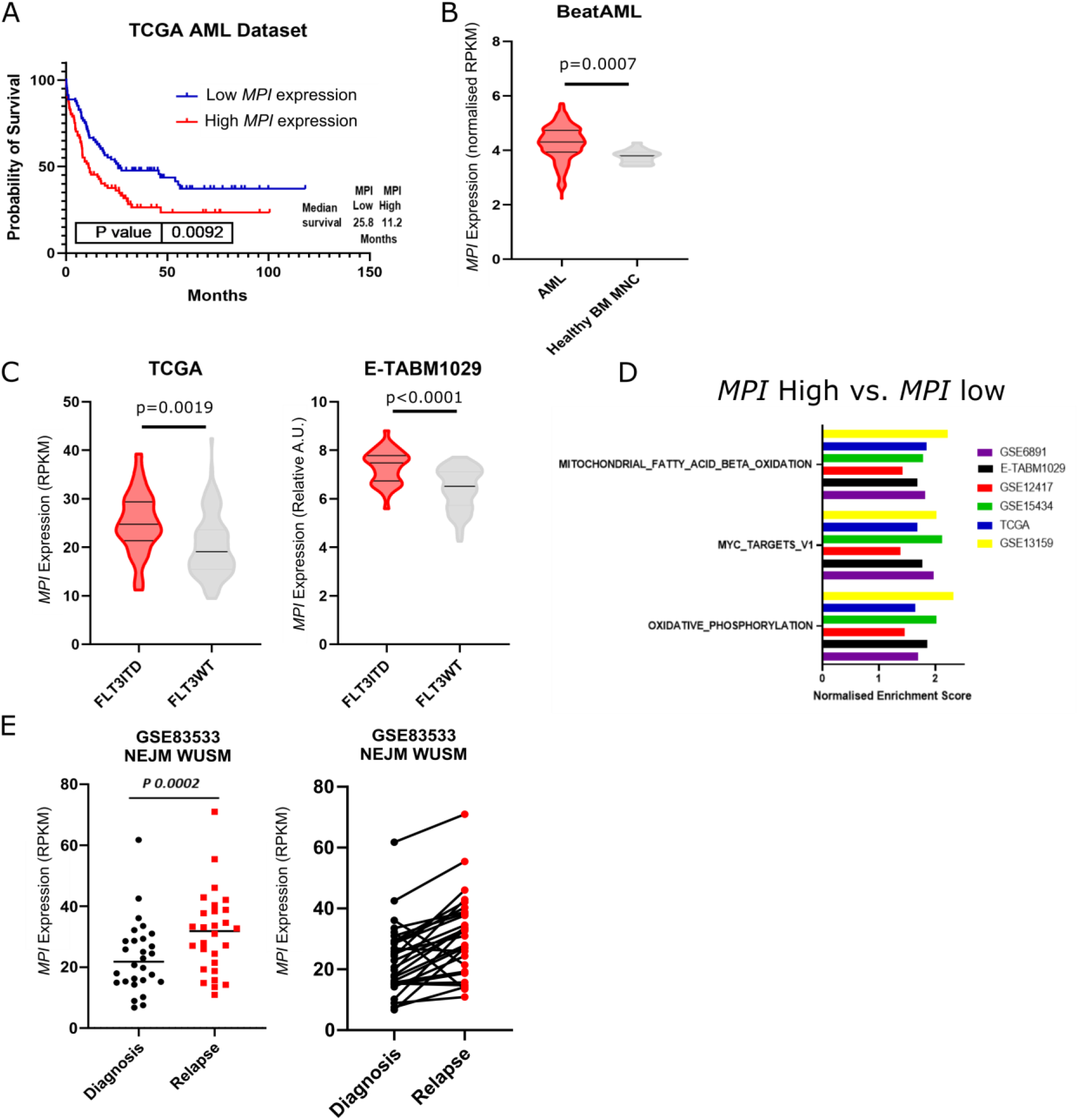
High MPI expression correlates with worse prognosis in AML and a gene signature associated with enhanced oxidative phosphorylation and fatty acid metabolism. **A** – Kaplan-Meier curve comparing survival of patients from the TCGA AML cohort separated by the top 50% and bottom 50% of *MPI* expression (data obtained from https://www.cbioportal.org/). Log-rank (Mantel-Cox) test; **B** – Violin plots of normalised *MPI* expression in AML samples compared to normal, healthy bone marrow mononuclear cells (MNC) from the BeatAML dataset (data obtained http://www.vizome.org/). Unpaired t-test; **C** – Violin plots comparing *MPI* expression in FLT3^ITD^ and FLT3^WT^ AML samples from the TCGA dataset (left) and E-TABM1029 dataset (right). Unpaired t-test; **D** – 3 significantly enriched gene signatures in *MPI* high expressing samples in 7 AML datasets (GSE6891, E-TABM1029, GSE12417, GSE15434, TCGA, GSE10358 and GSE13159); **E** – Relative expression of *MPI* at diagnosis and relapse in paired samples from GSE83533 dataset and data from manuscript 10.1056/NEJMoa1808777 (NEJM WUSM). Paired t-test.

### MPI inhibition sensitizes both wild-type and FLT3^ITD^ mutant AML cells to novel targeted and standard therapies

To test the functional role of MPI in AML, we generated MPI KO and respective control non-targeting(NT)gRNA (hereafter Control) AML cells by CRISPR editing (Extended Data Fig.2A) and confirmed that MPI KO sensitized FLT3^ITD^ AML cells to FLT3-TKI (Fig.2A). Moreover, both FLT3^ITD^ and FLT3^WT^ MPI KO AML cells were sensitized to standard cytarabine (AraC) chemotherapy in competition assays (Fig.2B). These effects were due to both reduced viability and proliferation and were rescued by adding mannose to the media thus confirming the specificity of MPI KO (Fig.2C-D and Extended Data Fig.2B-C). LC/MS analysis confirmed that mannose levels were reduced in MPI KO cells (Extended Data Fig.2D). Similar phenotypic effects were observed using the MPI inhibitor MLS0315771 (hereafter MLS) ^28^ and following genetic silencing by shRNA (Fig.2E and Extended Data Fig.2E-G). The specificity of MLS was confirmed by the lack of further toxicity in MPI KO cells (Extended Data Fig.2H). Finally, in *in vivo* experiments, mice transplanted with MPI KO cells were more sensitive to AC220 and AraC resulting in both significant prolonged survival and reduction in leukemia burden (Fig.2F and Extended Data Fig. I-M). Taken together these results show that MPI plays a role in resistance to both FLT3-TKI and AraC in AML cells.

**Figure 2.**
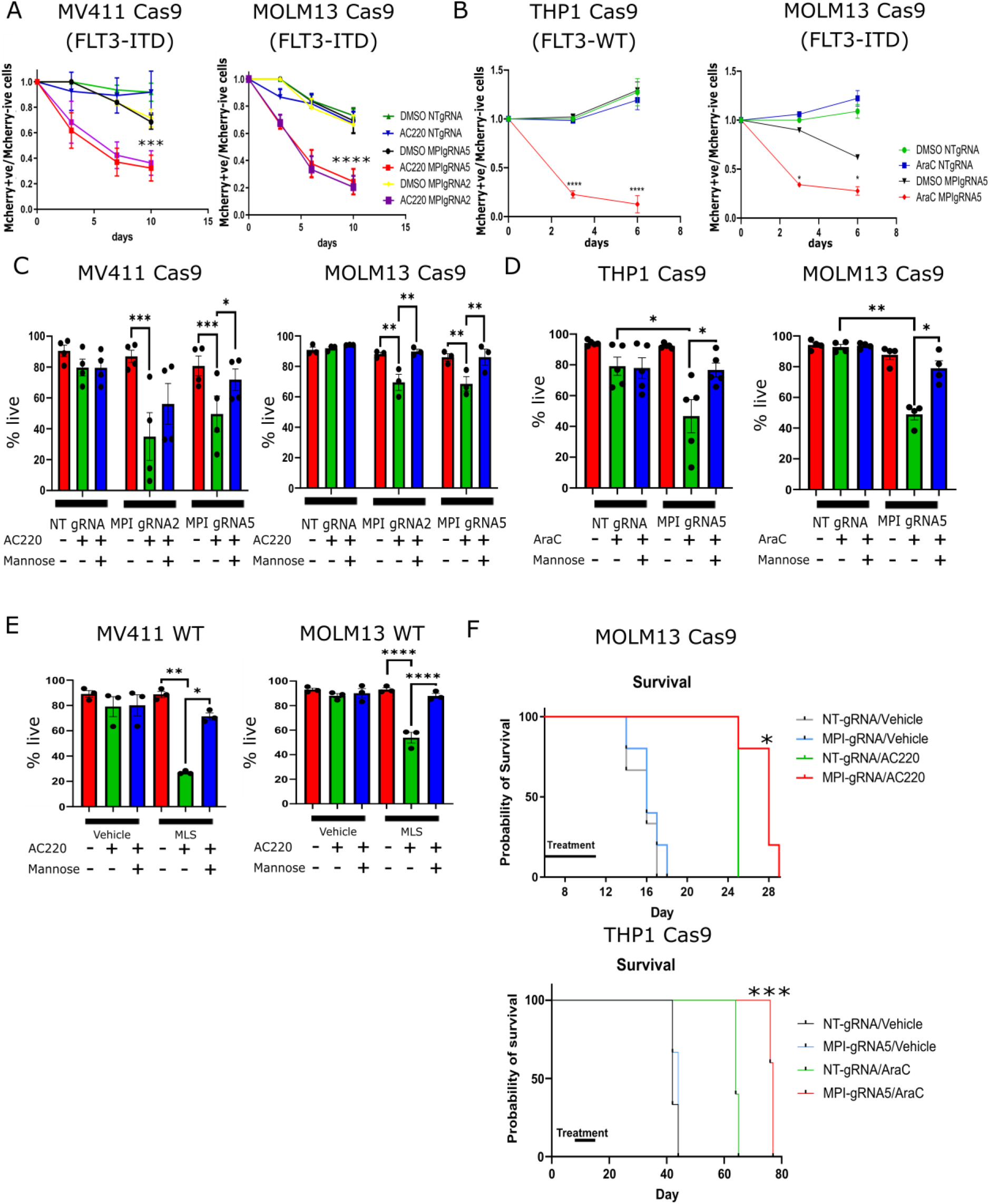
MPI inhibition sensitizes both wild-type and FLT3^ITD^ mutant AML cells to novel targeted and standard therapies. **A** – Competition growth assays measuring the ratio of mCherry positive to mCherry negative cells between WT MV411 (ie not expressing Cas9 or gRNAs and hence mCherry negative) and NT gRNA or MPI gRNA2 or gRNA5 (all mCherry positive) with and without AC220 (1nM) treatments (left) and WT MOLM13 (mCherry negative) and NT gRNA or MPI gRNA2 or gRNA5 (mCherry positive) with and without AC220 (1nM) treatments (right) over 10 days. N=3, 2 way Anova with Bonferroni correction for multiple comparisons; **B** - Competition growth assays measuring the ratio of mCherry positive to mCherry negative cells between WT THP1 (mCherry negative) and NT gRNA or MPI gRNA5 (mCherry positive) with and without AraC (1μM) treatments (left) and WT MOLM13 (mCherry negative) and NT gRNA or MPI gRNA5 (mCherry positive) with and without AraC (1μM) treatments (right) over 6 days. N=3, 2 way Anova with Bonferroni correction for multiple comparisons; **C** – Percentage of live cells NT gRNA, MPI gRNA2 and MPI gRNA5 MV411 (left) and MOLM13 (right) cells treated with vehicle (red), AC220 (1nM, green) or AC220 (1nM) and mannose (100μM, blue), as indicated. Treated for 6 days, N=4 for MV411, N=3 for MOLM13, 1 way Anova with Tukey’s correction for multiple comparisons; **D** – Percentage of live cells of NT gRNA and MPI gRNA5 THP1 (left) and MOLM13 (right) cells treated with vehicle (red), AraC (1μM, green) or AraC (1μM) and mannose (100μM, blue), as indicated. Treated for 3 days, N=4 for MOLM13, N=5 for THP1, 1 way Anova with Tukey’s correction for multiple comparisons; **E** - Percentage of live cells of WT MV411 (left) and WT MOLM13 (right) cells treated with vehicle, MLS0315771 (1μM), AC220 (1nM), mannose (100μM) or combinations of these as indicated. Treated for 6 days N=3, 1 way Anova with Tukey’s correction for multiple comparisons; **F** – Kaplan-Meier survival curve showing survival time of mice after transplantation with MOLM13 NT gRNA or MPI gRNA5 cells, with or without AC220 treatment (5mg/kg, top) and THP1 NT gRNA or MPI gRNA5 cells, with or without cytarabine treatment (50mg/kg, bottom), as shown by treatment bars. Log-rank (Mantel-Cox) test. For all panels, ns = not significant, *=p<0.05, **=p<0.01, ***=p<0.005, ****=p<0.001.

### MPI KO causes increased lipid uptake in AML cells

To understand how MPI inhibition increases sensitivity to AML therapies, we pursued an unbiased complementary metabolomic and transcriptional analysis (Supplemental Table 1 and 2). We focused on FLT3^ITD^ mutant cells treated with FLT3-TKI because MPI expression is higher in this AML subtype and adaptive resistance to FLT3-TKI is less well described thus necessitating a better understanding given the increasing clinical use of FLT3-TKI. Principal component analysis of both RNA-seq and untargeted metabolomics highlighted that MPI KO cells treated with FLT3-TKI separated from their respective Control and that the separation was partially rescued by addition of mannose to the media (Extended Data Fig.3A-B). We identified over 600 metabolites across different pathways through untargeted metabolomics. Unexpectedly, the most striking change detected was the significant increase in the intracellular levels of both long-chain saturated fatty acids (SFA) and polyunsaturated fatty acids (PUFA) in MPI KO cells treated with FLT3-TKI which was partially rescued by mannose supplementation (Fig.3A). Interestingly we also noted that both long and medium-chain acylcarnitines increase in AC220 treated MPI KO cells (Fig.3B) and this was coupled with a reduction in tricarboxylic acid (TCA) cycle intermediates in the same conditions (Extended Data Fig.3C). This metabolomic pattern suggests both an increase in uptake of long-chain fatty acids and defective mitochondrial fatty acid oxidation (FAO) in treated MPI KO cells. To specifically test for fatty acid uptake, we labelled the cells with the saturated lipid probe C1-Bodipy 500/510 C12 and showed that MPI KO cells significantly increased fatty acid uptake (Fig.3C and Extended Data Fig.3D). To clarify if there was a preferential uptake of SFAs, monounsaturated fatty acids (MUFAs) or PUFAs, we used neutral Bodipy 493/503 to stain cells fed specifically palmitate (16:0 SFA), oleate (18:1 MUFA) and arachidonic acid (20:4 PUFA). While we observed significant increase in uptake of all subtypes of fatty acids in MPI KO cells, arachidonic acid uptake appeared to be particularly enhanced in the KO cells and was also rescued by mannose (Fig.3D). This finding was corroborated by the metabolomic analysis as treated MPI KO cells showed higher increase in their PUFA and PUFA containing lipid species levels compared to MUFA and SFA (Extended Data Fig.3E). Interestingly CD36, a fatty acid transporter with prognostic significance in AML ^13,29^, was transcriptionally upregulated in MPI KO cells (Extended Data Fig.3F). To test if CD36 played a role in lipid uptake, we used its irreversible inhibitor sulfosuccinimidyl oleate (SSO) ^30^ and observed that, while CD36 inhibition reduced arachidonic acid uptake in MPI KO cells, it did not affect the uptake of the saturated lipid probe C1-Bodipy 500/510 C12 (Extended Data Fig.3G-H). Together these data are consistent with MPI KO cells having an increased lipid uptake which, at least for PUFAs, is driven by increased CD36 expression. Interestingly CD36 is known to preferentially favour uptake of long chain PUFAs ^31^.

**Figure 3.**
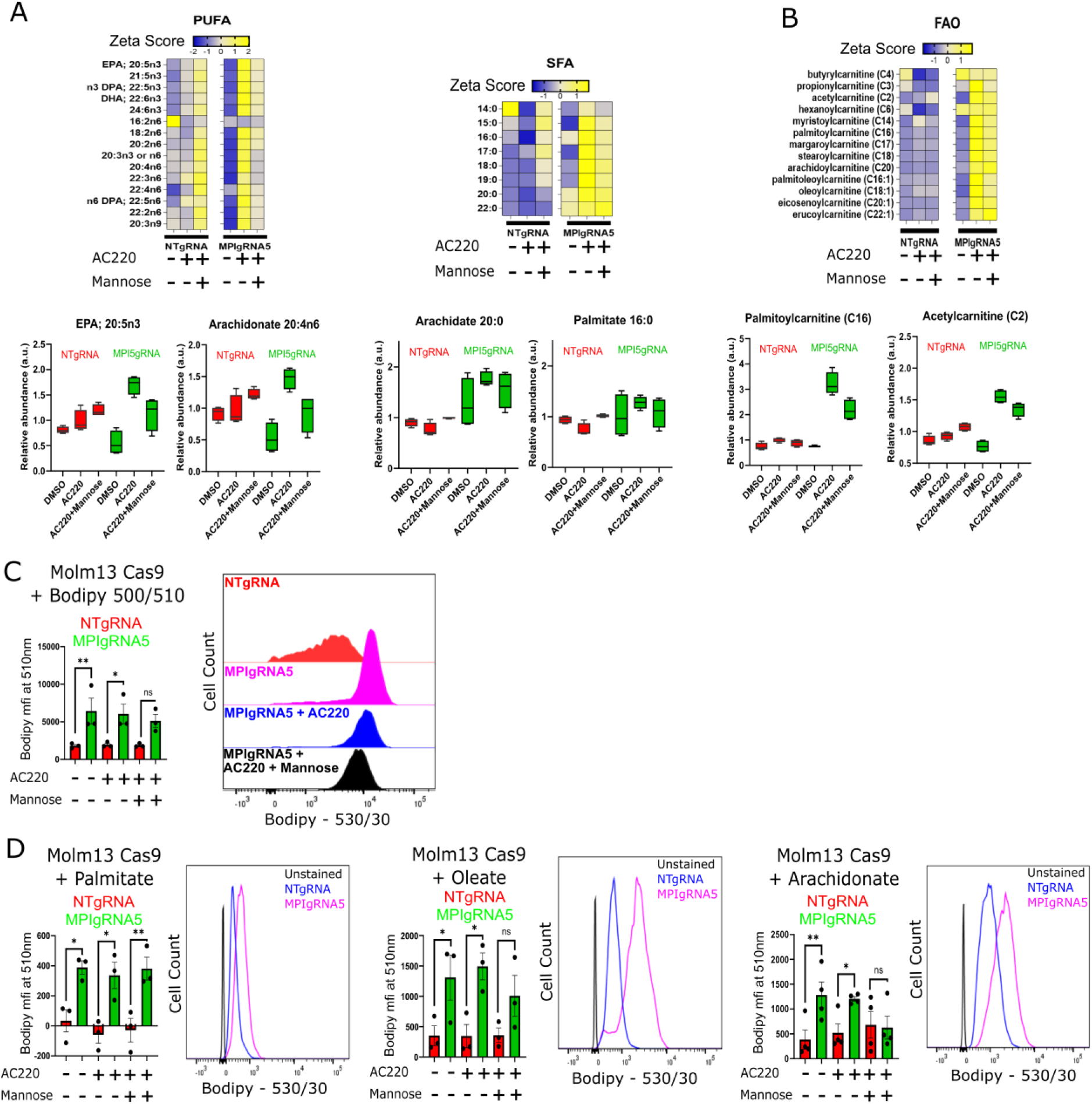
MPI KO causes increased lipid uptake in AML cells. **A** – Heat maps of Z-scores for relative changes of long chain polyunsaturated fatty acids (PUFA - n3, n6 and n9, left) and long chain saturated fatty acids (SFA, right), with box plots of selected lipids of each type below the heat maps from global metabolomics of MOLM13 NT gRNA or MPI gRNA5 cells treated with vehicle, AC220 (1nM) or AC220 (1nM) and mannose (100μM) as indicated for 48 hours; **B** – Heat maps of Z-scores for relative changes of fatty acid oxidation metabolism related fatty acid species, with box plots of selected lipids below the heat maps from global metabolomics of MOLM13 NT gRNA or MPI gRNA5 cells treated with vehicle, AC220 (1nM) or AC220 (1nM) and mannose (100μM) as indicated for 48 hours; **C**- Uptake of fluorescent C1-Bodipy 500/510 C12 by NT gRNA or MPI gRNA5 MOLM13 cells treated with vehicle, AC220 (1nM) or AC220 (1nM) and mannose (100μM) for 24 hours, with a representative flow cytometry plot (right). N=3, 1 way Anova with Tukey’s correction for multiple comparisons; **D** – Uptake of palmitate (50μM, left), oleate (50μM, middle) and arachidonic acid (1μM, right) by NT gRNA or MPI gRNA5 MOLM13 cells treated with vehicle, AC220 (1nM) or AC220 (1nM) and mannose (100μM) for 24 hours as indicated and measured by Bodipy 493/503 neutral lipid stain with representative flow cytometry plots comparing uptake by NT gRNA and MPI gRNA5 cells (with unstained sample, to the right of each graph). N=3 for palmitate and oleate, N=4 for arachidonic acid, 1 way Anova with Tukey’s correction for multiple comparisons. For all panels, ns = not significant, *=p<0.05, **=p<0.01, ***=p<0.005, ****=p<0.001.

### MPI KO results in gene expression and metabolic changes consistent with reduced FAO in AML cells

We then turned our attention to FAO, given our metabolomic profile was suggestive of a defective FAO. GSEA analysis of RNA-seq data demonstrated that oxidative phosphorylation and fatty acid metabolism signatures were upregulated in Control compared to MPI KO cells both in the absence and, even more, in the presence of AC220. Similar genesets were enriched in AC220 treated MPI KO cells supplemented with mannose when compared to AC220 treated MPI KO cells (Fig.4A and Extended Data Fig.4A). Consistent with this, *MPI* expression levels were positively correlated with genes involved in fatty acid metabolism across both TCGA and BeatAML datasets (Extended Data Fig.4B). Moreover, RT-QPCR showed reduction in expression levels of *CPT1A* and *PPARα*, a master regulator of lipid catabolism, in MPI KO compared to Control cells (Extended Data Fig.4C). Therefore, gene expression analysis suggest that treated MPI KO cells have defective FAO likely contributing to intracellular accumulation of both acylcarnitines and upstream fatty acids observed in metabolomic profiling.

**Figure 4.**
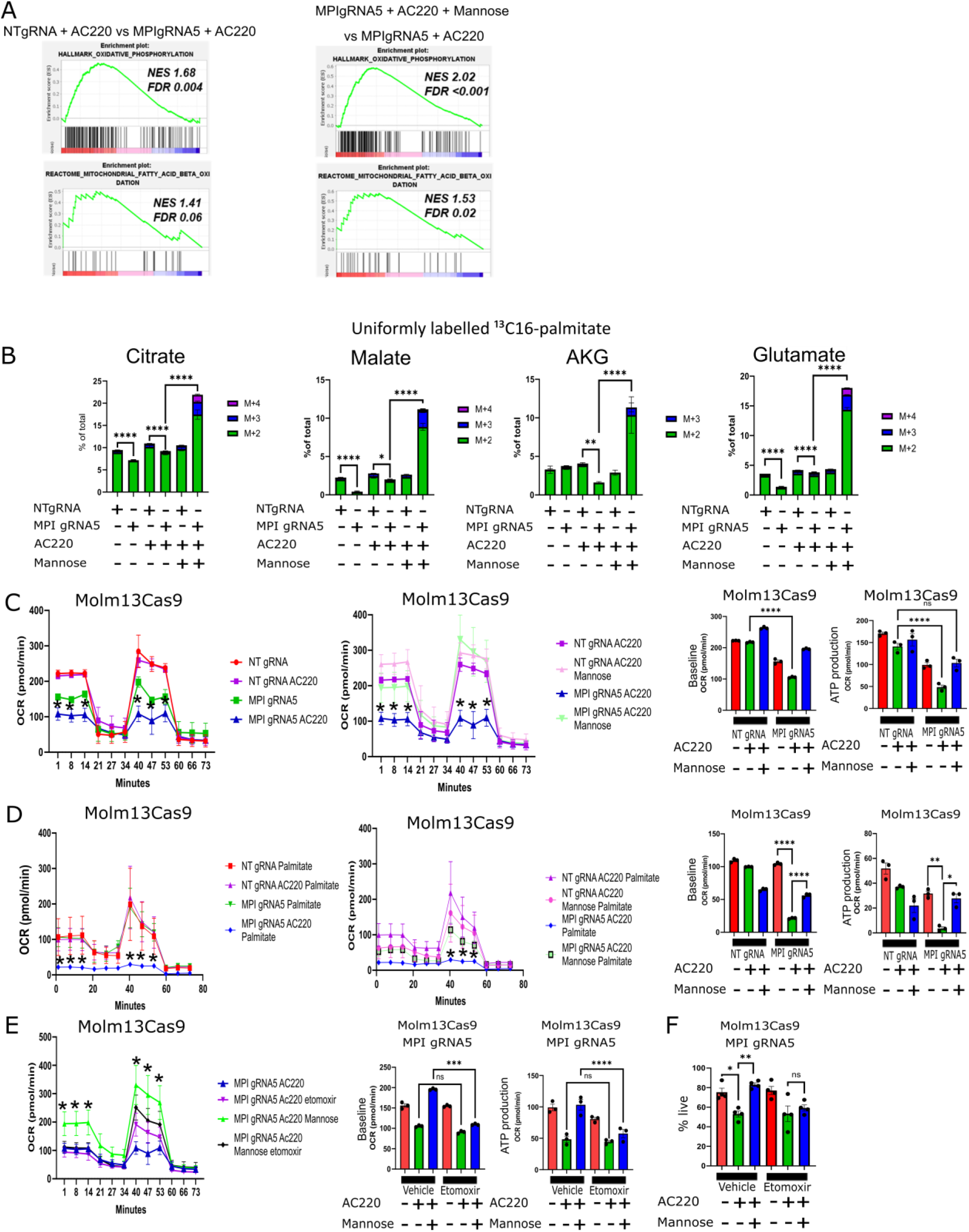
MPI KO results in gene expression and metabolic changes consistent with reduced FAO in AML cells. **A** – GSEA for Oxidative phosphorylation and Fatty acid oxidation gene signatures from RNA sequencing data comparing MOLM13 NT gRNA and MPI gRNA5 treated with AC220 (1nM, left) and MOLM13 MPI gRNA5 treated with AC220 (1nM) or AC220 (1nM) and mannose (100μM, right), FDR and NES from 1000 permutations; **B** – Percentage of TCA cycle intermediates and associated metabolites labelled with ^13^C from uniformly labelled ^13^C_16_-palmitate, Citrate (left), Malate (2^nd^ left), Alpha-ketoglutarate (AKG - 2^nd^ right) and Glutamate (far right) in MPI KO and NT MOLM13 cells treated with AC220 (1nm) and mannose (100μM) as indicated for 24 hours along with 50μM i^13^C_16_-palmitate. M+2 has 2 labelled C, M+3 has 3 labelled C, M+4 has 4 labelled C. N=5, 1-way anova with Tukey’s correction for multiple comparisons; **C** - SEAHORSE MitoStress tests showing oxygen consumption rate over time comparing NT gRNA and MPI gRNA5 MOLM13 cells treated with vehicle, AC220 (1nM) or AC220 (1nM) and mannose (100μM) as indicated after 72 hours of treatment, N=3, 2 way Anova with Sidak’s correction for multiple comparisons (two left panels). Baseline OCR and ATP production of NT gRNA and MPI gRNA5 MOLM13 cells treated with vehicle, AC220 (1nM) or AC220 (1nM) and mannose (100μM) as indicated after 72 hours of treatment, N=3, 1 way Anova with Tukey’s correction for multiple comparisons (2 right panels); **D** - SEAHORSE MitoStress tests showing oxygen consumption rate over time comparing MPI gRNA5 cells cultured overnight in substrate limited RPMI media without FBS and glutamine treated with vehicle, AC220 (1nM), mannose (100μM) or palmitate (50μM) in combinations as indicated N=3, 2 way Anova with Sidak’s correction for multiple comparisons (two left panels). Baseline OCR and ATP production of NT gRNA and MPI gRNA5 cells treated with palmitate (50μM) and vehicle, AC220 (1nM), mannose (100μM) in combinations as indicated, N=3, 1 way Anova with Tukey’s correction for multiple comparisons (2 right panels); **E** - SEAHORSE MitoStress tests showing oxygen consumption rate over time comparing MPI gRNA5 cells treated with AC220 (1nM), mannose (100μM) after 72 hours in combinations as indicated with or without etomoxir (50μM), N=3, 2 way Anova with Sidak’s correction for multiple comparisons (left panel). Baseline OCR and ATP production of MPI gRNA5 cells treated with vehicle, AC220 (1nM), mannose (100μM) after 72 hours in combinations as indicated with or without etomoxir (50μM). N=3, 1 way Anova with Tukey’s correction for multiple comparisons (2 right panels); **F** - Percentage of live MPI gRNA5 MOLM13 cells treated with vehicle, etomoxir (10μM), AC220 (1nM), mannose (100μM) or in combinations as indicated 6 days after treatment. N=4, 1 way Anova with Tukey’s correction for multiple comparisons. For all panels, ns = not significant, *=p<0.05, **=p<0.01, ***=p<0.005, ****=p<0.001.

To confirm reduction in FAO in MPI deficient cells, we grew cells in U-^13^C_16_-palmitate and found reduced palmitate labelling of several TCA intermediates in MPI KO cells, a phenotype rescued by mannose (Fig.4B and Extended Data Fig.4D). Real-time metabolic flux analysis also showed that MPI KO cells had reduced oxygen consumption rate (OCR) and ATP-linked OCR compared to Control, with the OCR further reduced in the presence of AC220 and rescued in the presence of mannose (Fig.4C). Similar findings were observed in THP1 cells treated with AraC (Extended Data Fig.4F). To specifically assess FAO we measured OCR using substrate limiting media in the presence of only palmitate. We first observed that with palmitate as the sole substrate, mannose supplementation rescued the respiration defect in treated MPI KO cells suggesting that MPI KO cells can engage efficiently in FAO only when supplemented with mannose (Extended Data Fig.4E). Moreover, in substrate limiting media with palmitate, both basal respiration and ATP production were significantly reduced in KO cells compared to Control following AC220 treatment with the effects rescued by mannose (Fig.4D). These data suggest MPI KO cells do not use palmitate efficiently as a source of energy, particularly in the presence of AC220, with this phenotype reversed by mannose. Indeed adding the CPT1A and FAO inhibitor etomoxir^11^ to AC220 or AraC did not further reduce basal respiration or ATP production in MPI KO cells. However, the re-addition of mannose resensitized MPI KO cells to the effects of etomoxir on OCR (Fig.4E and Extended Data Fig.4G). These effects correlated with changes in viability in response to etomoxir in MPI KO cells, as no reduction in viability beyond the effects of AC220 was observed in response to etomoxir, unless cells were cultured in the presence of mannose (Fig.4F). Conversely viability of AC220 or AraC treated MPI KO was rescued by the PPARα agonist and FAO activator fenofibrate^32^ to a degree similar to mannose (Extended Data Fig.4H). Finally we did not rescue viability of treated MPI KO cells using the cell permeable TCA cycle intermediate dimethyl-alpha-ketoglutarate (DMaKG) (Extended Data Fig.4I). These data suggest that MPI KO cells have defective FAO but the effects on viability of MPI KO are driven by both defective FAO and enhanced lipid uptake causing an increase in intracellular fatty acid levels rather than simply a reduction in TCA cycle activity and oxidative phosphorylation.

### MPI KO cells activate the ATF6 arm of the unfolded protein response (UPR) to inhibit FAO in AML cells

MPI deficiency leads to a defect in protein glycosylation^17^ and as predicted MPI KO cells showed reduced protein glycosylation as confirmed by their lower lectin staining. This was also associated with concurrent accumulation of misfolded protein (Extended Data Fig.5A-B). These effects correlated, as expected, with upregulation of UPR genes in MPI KO cells although in particular the ATF6 arm of the UPR, which is particularly sensitive to changes in glycosylation^33^, was upregulated (Fig.5A). To validate this, we transduced cells with an ATF6-YFP reporter lentivirus (Extended Data Fig.5C)^34^ and observed significant ATF6 activation in MPI KO cells which was reversed by mannose (Fig.5B). We then used immunofluorescence to track localisation of ATF6 in MPI KO cells and showed that ATF6 localised in the nucleus of MPI KO cells and this was reversed by mannose (Fig.5C-D and Extended Data Fig.5D). Similarly western blot showed higher levels of cleaved ATF6 consistent with its activation in MPI KO cells compared to Control cells which was rescued by mannose (Extended Data Fig.5E). Interestingly both in the ATF6 reporter assay and by western blot, AC220 treatment appeared also to reduce ATF6 activation although this was not seen by immunofluorescence. This discrepancy might reflect sensitivity of the assay and/or be related to the effects of AC220 on protein translation/ER load and ATF6 activation at the assessed timepoint. Conversely we did not observe consistent significant changes in the activation of the other branches of the UPR in MPI KO cells (Extended Data Fig.5F). To functionally test the effects of ATF6 activation in MPI KO cells, we used the highly specific ATF6 inhibitor Ceapin A7 which retains ATF6 in the endoplasmic reticulum thus preventing its nuclear translocation and ability to activate transcription of its target genes^35,36^. Treatment with Ceapin A7 phenocopied mannose supplementation by reversing ATF6 cleavage and nuclear localisation (Fig.5C-D and Extended Data Fig.5G). Ceapin A7 also rescued viability and OCR of MPI KO cells treated with AC220 to the same degree of mannose and resensitized MPI KO cells to the effects of etomoxir on OCR (Fig.5E-F). Similar effects were observed following silencing of ATF6 (Extended Data Fig.5H-I). Conversely both the UPR activator with reported preferential activity on ATF6 AA147^37^ (Extended Data Fig.5M) and the canonical glycosylation inhibitor and UPR activator tunicamycin mimicked the effects of MPI KO on cell viability and prevented mannose rescue (Extended Data Fig.5J-K). Finally inhibitors of the other UPR branches did not affect viability of treated MPI KO cells (Extended Data Fig.5L). Therefore despite the fact that the UPR response is often non-selective and there is evidence that ATF6 can lead to activation of other arms of the UPR response^38,39^, we observe in our system a preferential activation of ATF6. This might be due to low level and chronic endoplasmic reticulum stress activating preferentially the ATF6 arm of the UPR. Interestingly this is consistent with previous literature showing that ATF6 is able to refold proteins and prevent further stress and activation of the other arms of UPR which happens only when the refolding ability driven by ATF6 is overwhelmed^38,40^. Activation of the ATF6 arm of the UPR has been reported to inhibit the transcription/activity of the master regulator of lipid catabolism PPARα ^41^ and interestingly both Ceapin A7 and AA147, beside modulating the expression of canonical ATF6 targets *ERO1B* and *HERPUD1* as expected, respectively increased and decreased the expression of *PPARα* and *CPT1A*, the rate limiting step in FAO, thus phenocopying the transcriptional consequences of MPI KO and mannose rescue (Extended Data Fig.5M-N). Interestingly, *MPI* expression levels were negatively correlated with *ERO1B* and *HERPUD1* expression across both TCGA and BeatAML datasets (Extended Data Fig.5O). Taken together these data support a model where MPI KO cells preferentially activate the ATF6 arm of the UPR leading to reduced FAO.

**Figure 5.**
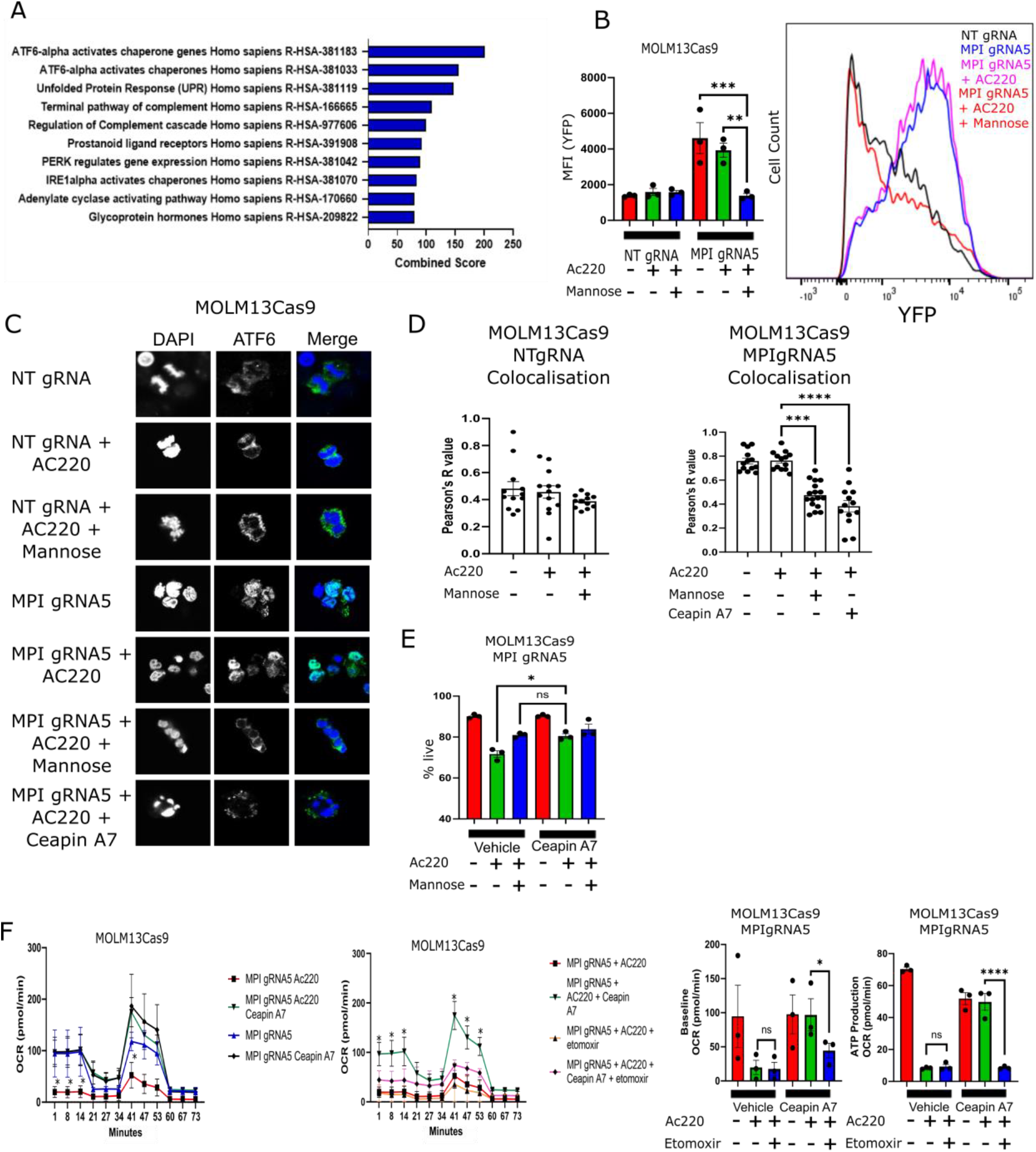
MPI KO cells activate the ATF6 arm of the unfolded protein response (UPR) to inhibit oxidative phosphorylation metabolism in AML cells. **A** – Top combined score gene signatures from RNA sequencing enriched in MPI gRNA5 MOLM13 cells compared to NT gRNA MOLM13 cells. Analysis performed with Enrichr; **B** – Mean fluorescence intensity of YFP from MOLM13 NT gRNA or MPI gRNA5 cells expressing a YFP linked ATF6 reporter treated with vehicle, AC220 (1nM), mannose (100μM) or combinations as indicated (left panel) with a representative flow cytometry plot (left right panel). N=3, 1 way Anova with Tukey’s correction for multiple comparisons; **C** – Confocal microscopy images of NT gRNA and MPI gRNA5 MOLM13 cells stained for DAPI (blue, 1^st^ column) and ATF6 (green, 2^nd^ column) with a merged image (3^rd^ column), treated with vehicle, AC220 (1nM), mannose (100μM), Ceapin A7 (1μM) or in combinations as indicated 72 hours after treatment; **D** - Colocalisation analysis from immunofluorescence images of NTgRNA (left) and MPIgRNA5 (right) MOLM13 cells treated with vehicle, AC220 (1nM), mannose (100μM) and Ceapin A7 (1μM) in combinations as indicated. Analysis performed with Coloc2 plugin in ImageJ, ordinary 1 way Anova with Tukey’s correction for multiple comparisons; **E** – Percentage of live cells of MPI gRNA5 MOLM13 cells treated with vehicle, AC220 (1nM), mannose (100μM), Ceapin A7 (1μM) or in combinations as indicated 72 hours after treatment. N=3, 1 way Anova with Tukey’s correction for multiple comparisons; **F** - SEAHORSE MitoStress assay comparison of MPI gRNA5 MOLM13 cells, treated with AC220 (1nM), Ceapin A7 (1μM), etomoxir (10μM) or in combinations as indicated 72 hours after treatment. N=3, 2 way Anova with Sidak’s correction for multiple comparisons, with MPI gRNA5 AC220 as control population (2 left panels). Baseline OCR and ATP production of NT gRNA and MPI gRNA5 cells treated with vehicle, AC220 (1nM), Ceapin A7 (1μM), etomoxir (50μM) or in combinations as indicated, N=3, ns = not significant, 1 way Anova with Tukey’s correction for multiple comparisons (2 right panels). For all panels, ns = not significant, *=p<0.05, **=p<0.01, ***=p<0.005, ****=p<0.001.

### MPI KO drives lipid peroxidation and ferroptotic cell death in AML cells

The metabolic phenotype of MPI KO cells with accumulation of PUFA and reduced FAO is reminiscent of that observed in cancer cells prone to ferroptotic cell death, i.e. clear cell renal carcinoma^42,43^. Moreover MPI KO cells had lower levels of intracellular cysteine and higher levels of 4-hydroxy-nonenal-glutathione and ophthalmate (Fig.6A) markers of oxidative stress and lipid peroxidation^44,45^ which was further increased by AC220 a known driver of oxidative stress in these cells^9^. Interestingly a ferroptotic gene signature^46^ was upregulated in treated MPI KO cells compared to both Control and MPI KO mannose treated cells (Fig.6B). These observations led us to test if cell death of treated MPI KO cells was driven by lipid peroxidation and ferroptosis. Indeed Bodipy 581/591 C11 staining of MPI KO and knock-down (KD) cells confirmed higher levels of lipid peroxidation in treated MPI KO/KD cells, that was rescued by mannose (Fig.6C and Extended Data Fig.6A-B). Moreover, cell death and induction of lipid peroxidation in both AC220 and AraC treated MPI KO cells was rescued by the radical-trapping antioxidant ferrostatin and liproxstatin, known anti-ferroptosis agents^47^ (Fig.6D and Extended Data Fig.6C-E). Interestingly Ceapin A7 and AA147 respectively mimicked and inhibited the effects of mannose on lipid peroxidation supporting the role of ATF6 activation in driving the ferroptotic phenotype (Extended Data Fig.6F-G). Conversely neither PERK or IRE1 inhibition had significant effects on lipid peroxidation (Extended Data Fig. 6H-I). MPI KO cells were more sensitive to PUFA (arachidonic acid) toxicity which is rescued by mannose and SSO thus reinforcing the role of CD36 in PUFAs uptake (Extended Data Fig.7A-B). Conversely growing MPI KO cells in delipidated media reduced their sensitivity to AC220 (Extended Data Fig.7C). However SSO did not affect lipid peroxidation (Extended Data Fig. 7D) probably because it affects uptake of only some fatty acid species (Extended Data Fig.3G-H) and has no effects on FAO. This also suggests that sensitivity to arachidonic acid toxicity is not just due to increased lipid peroxidation. MPI KO cells were also more sensitive to the ferroptotic inducing agents erastin and RSL3 (Extended Data Fig.7E). Interestingly, and consistent with the reduced cysteine levels in MPI KO cells, surface expression of the cysteine transporter and erastin target SLC7A11/xCT was downregulated in MPI KO cells further explaining their tendency to undergo ferroptosis (Extended Data Fig.7F). We also did not detect significant upregulation of cleaved PARP in treated MPI KO cells and neither the apoptosis/pan caspase inhibitor Z-VAD or necroptosis inhibitor Necrostatin-1 rescued cell death in both AC220 and AraC treated MPI KO cells (Extended Data Fig.7G-H). Finally to validate the role of ferroptosis induction in driving cell death *in vivo* mice transplanted with THP1 Control and MPI KO cells were treated with AraC +/-liproxstatin. As expected liproxstatin treatment reduced disease latency in AraC treated mice transplanted with MPI KO cells and this correlated with reduced staining of the leukemic cells for 4-HNE, a marker of lipid peroxidation, *in vivo* (Fig 6E-F and Extended Data Fig. 7I-J). Overall these data show that, following chemo/targeted therapy treatment the metabolic rewiring caused by MPI KO primes the cells to lipid peroxidation and ferroptotic cell death.

**Figure 6.**
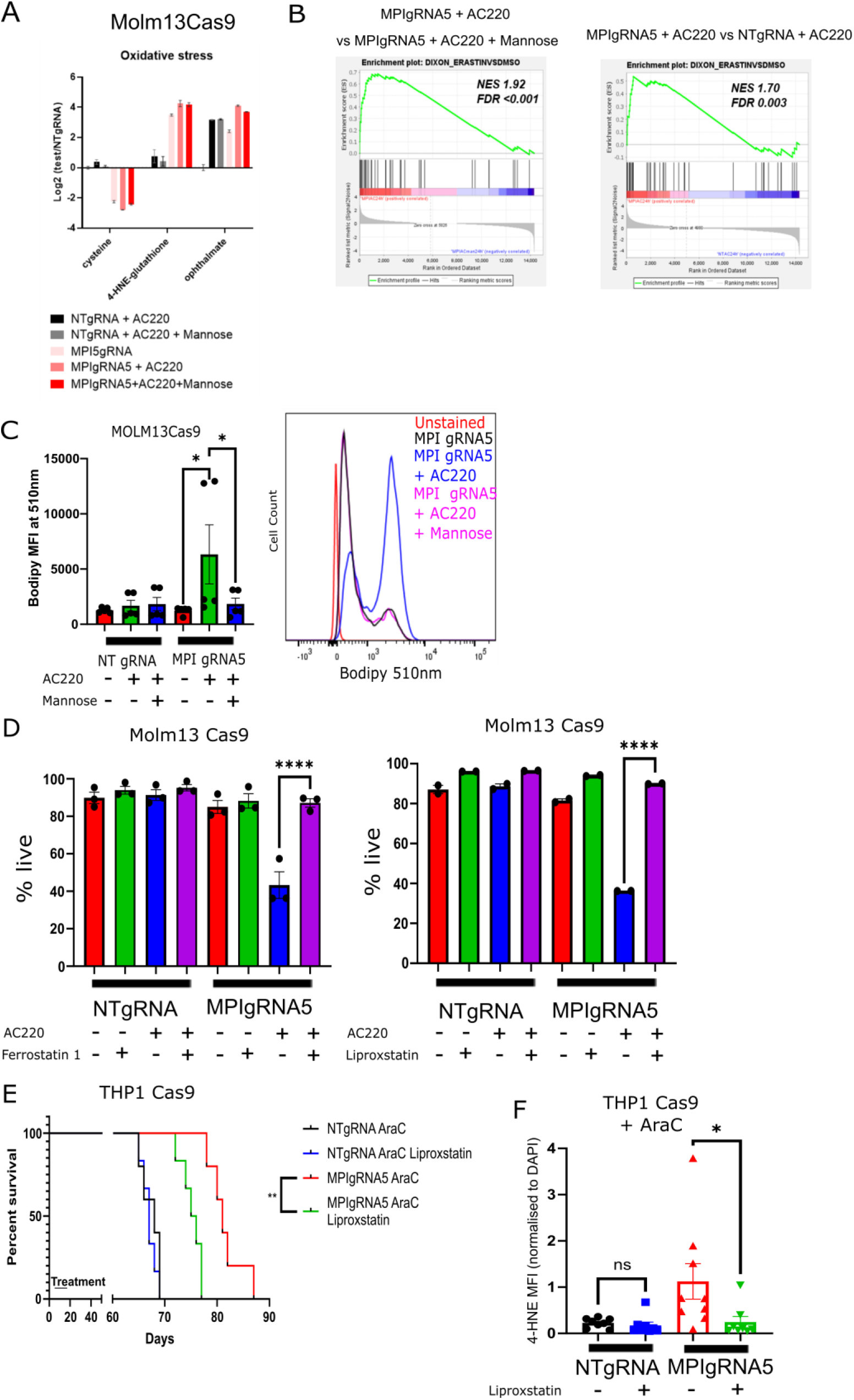
MPI KO drives lipid peroxidation and ferroptotic cell death in AML cells. **A** – Levels of ROS and lipid peroxidation linked metabolites from global metabolomics profiling of NT gRNA and MPI gRNA5 treated with vehicle, AC220 (1nM), mannose (100μM) or in combinations as indicated for 48 hours; **B** – GSEA for an erastin induced (ferroptotic) gene signature^43^ from RNA-seq comparing MPI gRNA5 with AC220 (1nM) to MPI gRNA5 with AC220 (1nM) and mannose (100μM) (left) and MPI gRNA5 with AC220 to NT gRNA with AC220 (right). FDR and NES from 1000 permutations; **C** – Mean fluorescence intensity at 510nm of Bodipy 581/591 C11, which shows level of lipid peroxidation, in NT gRNA and MPI gRNA5 MOLM13 cells treated with AC220 (1nM), mannose (100μM) or in combinations as indicated 72 hours after treatment. N=3, 1 way Anova with Tukey’s correction for multiple comparisons (left). Representative plots of flow cytometry fluorescence intensity plot of MPI gRNA5 with AC220 (1nM), mannose (100μM) or in combinations as indicated along with an unstained MPI gRNA5 control at 510nm (right); **D** – Percentage of live NT gRNA and MPI gRNA5 MOLM13 cells treated with vehicle, AC220 (1nM), mannose (100μM), ferrostatin-1 (5μM, left), liproxstatin (1μM, right) or combinations as indicated for 6 days. N=3 for ferrostatin, N=2 for liproxstatin, 1 way Anova with Tukey’s correction for multiple comparisons. **E** - Kaplan-Meier survival curve showing survival time of mice after transplantation with THP1 NT gRNA or MPI gRNA5 cells, treated with cytarabine (50mg/kg) for 7 days concurrent with or without liproxstatin treatment (10mg/kg) for 10 days. Log-rank (Mantel-Cox) test; **F** – Mean fluorescence intensity of 4-hydroxynonenal normalised to DAPI fluorescence from immunofluorescence images of THP-1 cells sorted from mice after transplantation with THP1 NT gRNA or MPI gRNA5 cells and soon after treatment with cytarabine (50mg/kg) for 7 days concurrent with or without liproxstatin treatment (10mg/kg) for 10 days. 1 way Anova with Tukey’s correction for multiple comparisons. For all panels, ns = not significant, *=p<0.05, **=p<0.01, ***=p<0.005, ****=p<0.001.

### MPI depletion in primary FLT3^ITD^ AML samples causes ATF6 activation, lipid peroxidation and sensitization to FLT3-TKI therapy

We finally validated the effects of MPI depletion in primary FLT3^ITD^ AML samples (Supplemental Table 3). Following MPI KD (Extended Data Fig.8A-B), FLT3^ITD^ primary AML MNC were sensitized to FLT3-TKI therapy and this could be rescued by mannose (Fig.7A). Conversely normal CD34^+^ cells were not sensitive to MPI KD both in the absence or presence of AC220 (Fig.7B). Consistent with observations in AML cell lines, MPI KD primed FLT3^ITD^ AML MNC to lipid peroxidation following FLT3-TKI therapy and led to higher levels of nuclear ATF6, with both effects rescued by the addition of mannose (Fig.7C-D). Conversely a smaller induction of lipid peroxidation was observed in normal CD34^+^ cells upon MPI KD which was not enhanced by FLT3-TKI or rescued by mannose (Extended Data Fig.8C). Moreover the viability of primary MPI KD FLT3^ITD^ AML MNC treated with AC220 was rescued by ferrostatin and liproxstatin confirming that the mechanism of death in primary AML MNC was also due to ferroptosis (Fig.7E). These experiments validate the phenotype driven by MPI inhibition in primary AML MNC while demonstrating that MPI KD is tolerated by normal hematopoietic stem/progenitor cells (HSPC).

**Figure 7.**
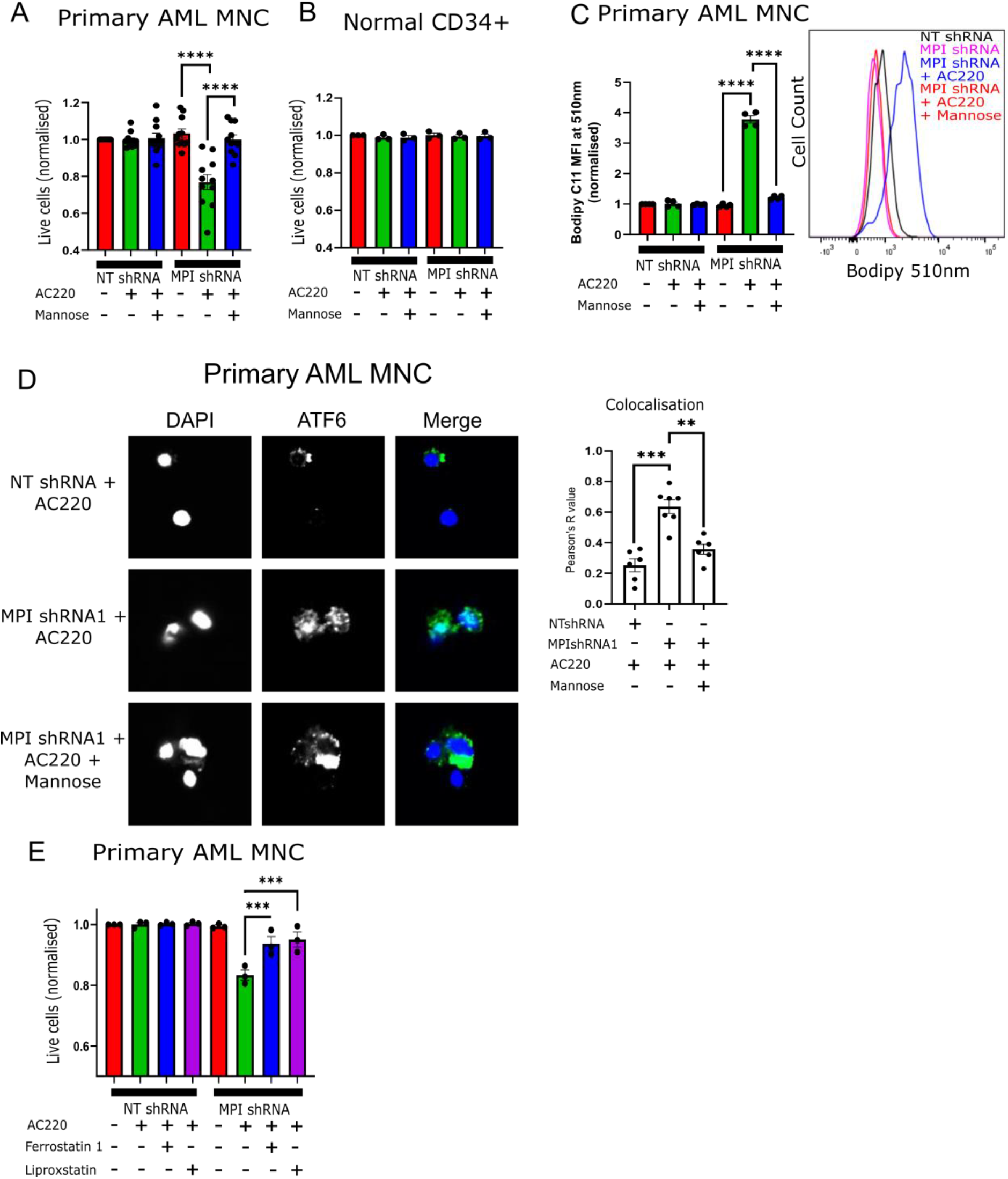
MPI depletion in primary FLT3^IT^D AML samples causes ATF6 activation, lipid peroxidation and sensitization to FLT3-TKI therapy. **A** - Percentage of live cells of NT shRNA and MPI shRNA1 primary FLT3^ITD^ AML MNC treated with vehicle, AC220 (2.5nM) or AC220 (2.5nM) and mannose (100μM), as indicated normalised to NT shRNA with vehicle for each sample. Treated for 3 days, N=11, 1 way Anova with Tukey’s correction for multiple comparisons; **B** - Percentage of live cells of NT shRNA and MPI shRNA1 normal CD34+ primary cells treated with vehicle, AC220 (2.5nM) or AC220 (2.5nM) and mannose (100μM), as indicated normalised to NT shRNA with vehicle for each sample. Treated for 3 days, N=3; **C** - Mean fluorescence intensity at 510nm of Bodipy 581/591 C11, which shows level of lipid peroxidation, in NT shRNA and MPI shRNA1 FLT3^ITD^ primary AML MNC treated with vehicle, AC220 (2.5nM) or AC220 (2.5nM) and mannose (100μM), as indicated normalised to NT shRNA with vehicle at 72 hours after treatment. N=4, 1 way Anova with Tukey’s correction for multiple comparisons (left). Representative plots of flow cytometry fluorescence intensity plot NT shRNA and MPI shRNA1 with AC220 (2.5nM) and mannose (100μM) as indicated at 510nm (right); **D** - Confocal microscopy images of FLT3^ITD^ primary AML MNC stained for DAPI (blue, 1^st^ column) and ATF6 (green, 2^nd^ column) with a merged image (3^rd^ column), treated with vehicle, AC220 (2.5nM) and mannose (100μM) combinations as indicated 72 hours after treatment with colocalisation analysis (right graph), 1 way Anova with Tukey’s correction for multiple comparisons; **E** - Percentage of live cells of NT shRNA and MPI shRNA1 primary FLT3^ITD^ AML MNC treated with vehicle, AC220 (2.5nM), Ferrostatin-1 (5μM) or liproxstatin (1μM) as indicated, normalised to NT shRNA with vehicle for each sample. Treated for 3 days, N=3, 1 way Anova with Tukey’s correction for multiple comparisons. For all panels, ns = not significant, *=p<0.05, **=p<0.01, ***=p<0.005, ****=p<0.001.

## Discussion

Our data show for the first time that targeting MPI and MM sensitizes AML cells to AraC and FLT3-TKI. Recent reports, including one in leukemic cells^48^, suggest that MPI could be a therapeutic target in cancer therapy^22,23^. Most of these reports however suggest that MPI inhibition would limit the ability of cancer cells to withstand the toxicity of an exogenous high-dose mannose diet. Our data instead support the role of MPI, at physiological concentrations of mannose, as an enzyme important in supporting AML cells survival following therapy through its ability to modulate MM, protein glycosylation and activity of the ATF6 arm of the UPR. It is worth noting that the effects of MPI inhibition *in vivo* could be blunted by the endogenous mannose production and dietary mannose and this could partly limit the role of MPI as a therapeutic target. Conversely, given that glucose, which is 100 fold more abundant than mannose in human plasma^14^, is the biggest source of mannose within the cells^15^ and considering the higher expression levels of MPI in AML samples, it is plausible that MPI inhibition will be detrimental to AML cells survival and its effects not rescued sufficiently by the relatively low levels of mannose in plasma. This outcome would be expected particularly if associated with reduced glucose metabolism, such as when combined with FLT3-TKI. Interestingly congenital MPI defects in humans are successfully treated with mannose supplements suggesting that physiological levels of mannose are unable to overcome glycosylation defects in MPI deficient patients.

An ideal drug target requires a robust therapeutic window in order to be exploitable. Although the embryonic lethality of *Mpi* in animal models is a reason for concern, our preclinical data suggest that HSPC are less reliant on MPI activity than AML cells. Moreover the lack of hematopoietic defects in humans carrying MPI mutation (albeit in a hypomorphic state) suggest that MPI inhibition might be tolerable for HSPC^18^. Elevated mannose metabolism may be more critical for developing tissues, as suggested by a recent report using hypomorphic alleles of PMM2, the enzymatic step following MPI in MM, that demonstrated that, while mannose sugar donors are necessary for embryonic development, they are only needed at very low levels after development^49^. Thus, the activity of MPI might be crucial for rapidly dividing cells but less essential in adult established tissues, a finding not uncommon in drug development where often embryonically lethal/essential genes are shown to be good drug targets with acceptable toxicity in the adult/homeostatic context^50^.

MPI validity as a therapeutic targets has been related to its ability to detoxify exogenous high levels of mannose^23,51^ and interestingly we also observed that feeding AML cells, particularly MPI KO, millimolar concentrations of mannose is toxic (not shown). However our mechanistic studies show that, even in the presence of physiological mannose concentrations, MPI inhibition can increase the toxicity of FLT3-TKI or AraC to AML cells due to defective fatty acid metabolism. While other reports on the role of MPI in cancer have established its role in supporting or rewiring central carbon/glucose metabolism, none had previously unveiled a connection between MM and fatty acid metabolism, although the ability of exogenous mannose to upregulate FAO has been observed in T cells^52^. We show that the effects on fatty acid metabolism are prompted by reduction in protein N-glycosylation, a known consequence of MPI depletion, which likely drives the preferential activation of the ATF6 arm of the UPR^33^. Interestingly reduced protein N-Glycosylation has already been shown to be a therapeutic vulnerability in solid cancers^22^ and leukemia^53^ and specifically in FLT3^ITD^ AML due to reduced surface FLT3 expression upon glycosylation inhibition^54^. While we could show a pattern of reduced protein glycosylation in our models, we did not detect consistent findings for reduced FLT3 expression (not shown) thus making this mechanism less likely. On the other hand, we observed that activation of the ATF6 arm of the UPR led to downregulation of FAO in AML MPI KO cells through transcriptional downregulation of PPARα and other key FAO genes. FAO has emerged as a therapeutic target in AML^11,55^ and is important in resistance to both standard^13^ and novel therapies^56^ via its ability to support oxidative phosphorylation. Therefore our findings support the role of FAO as a resistance mechanism in AML. Although FLT3-TKI and AraC have different mode of actions, resistance to both these drugs relies on enhanced FAO^13,57,58^ and likely explains why MPI KO by causing defective FAO and increased PUFAs levels leads to sensitization to both therapies. Moreover the role of reduced glycosylation and ATF6 activation in driving these effects further suggest that the main role of MPI in our systems is the production of mannose from glucose, rather than the opposite, and explains why MPI inhibition was able to cause cell death in the absence of high mannose concentration.

Although the effects of MPI KO on cell death could be partly driven by defective TCA cycle activity leading to ATP depletion, we did not rescue the effects of MPI KO by supplementing the TCA cycle intermediate DMaKG. Instead AML MPI KO cells, particularly in the presence of FLT3-TKI, displayed features consistent with a state primed for ferroptotic cell death. These included reduced FAO, increased CD36 expression and PUFA uptake, reduced expression of SLC7A11, the cysteine transporter and target of the ferroptotic inducing agent erastin^59^ and increased sensitivity to ferroptosis inducing agents and PUFA-induced toxicity. Ferroptosis is a non-apoptotic, iron dependent form of regulated cell death that is specifically characterised by the accumulation of lipid peroxides^60^. Susceptibility to ferroptosis has been shown in solid cancer models to be a feature of cells in a therapy resistant state because of their high dependency on the lipid hydroperoxides quenching proteins such as glutathione peroxidase 4^61^. Reduced FAO and downregulation of PPARα the master regulator of lipid catabolism have also been linked to susceptibility to ferroptosis in other cancer models^42,43,62^ while increased uptake of PUFA via CD36 has been shown to trigger ferroptotic cell death/susceptibility in T cells^63,64^. In this respect, it is worth noting that increased lipid uptake can enrich cancer cells membranes in PUFA since in contrast to de-novo synthetisized FA, diet and stroma derived FA are enriched for PUFA thus increasing susceptibility to ferroptosis ^65^. Moreover, there is evidence that PUFA are most readily beta-oxidised in mammalian cells compared to SFA ^66^. Therefore, a combination of increased lipid uptake and reduced FAO will lead to preferential accumulation of PUFA with an increased tendency towards lipid peroxidation, as seen in our MPI KO cells. Our rescue experiments with radical trapping agents confirm that ferroptosis is mostly driving cell death in MPI KO cells and future studies will be needed to further define the molecular mechanisms through which MPI KO drives increased PUFA uptake, CD36 expression and reduced cysteine uptake and SLC7A11 expression. Moreover, our findings also suggest that, similarly to persistent cells in solid cancers^61^, triggering ferroptosis can be leveraged to drive cell death in therapy-resistant AML cells. However, the role of ferroptosis susceptibility in AML therapy resistant/persister cells will require further studies.

In conclusion, our work supports the role of metabolic rewiring in driving therapy resistance in AML and demonstrates, for the first time, that targeting MPI and MM sensitizes AML cells to AraC and FLT3-TKI. Mechanistically we unveil a novel connection between MM and fatty acid metabolism, via preferential activation of the ATF6 arm of the UPR, leading to cellular PUFA accumulation, lipid peroxidation and ferroptotic cell death (Fig.8). Finally, our findings also suggest that triggering ferroptosis could be used therapeutically to eradicate therapy-resistant AML cells.

**Figure 8.**
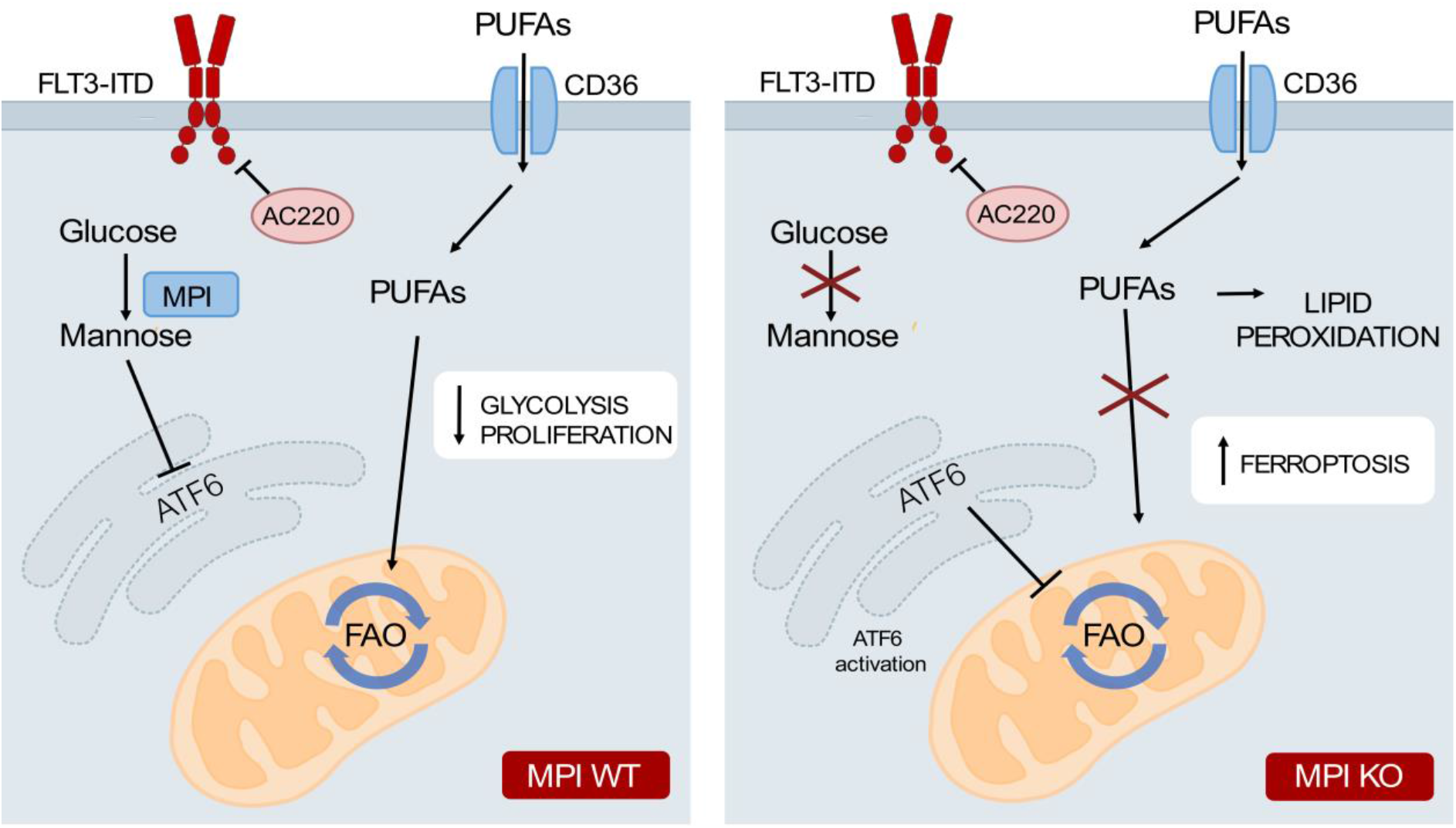
Model: Loss of MPI leads to cell death in AML through inhibition of FAO leading to PUFA accumulation and ferroptosis. A model of the proposed mechanism. WT AML cells treated with both standard and FLT3 inhibitor therapies are able to escape cell death by adapting their metabolism, in this case by switching from glycolysis to fatty acid oxidation. Conversely treated AML cells with depleted MPI have preferential activation of the ATF6 arm of the unfolded proteins response which inhibits fatty acid oxidation. This is paired with increased uptake of fatty acids, particularly polyunsaturated fatty acids, by MPI depleted cells. Both these effects lead to intracellular accumulation of PUFAs and PUFAs containing lipid species. These undergo lipid peroxidation leading to ferroptotic cell death in these cells.

## Acknowledgements

The authors wish to thank the Barts Cancer Institute tissue bank for sample collection and processing. This research was supported by the BCI Flow cytometry facility (CRUK Core Award C_16_420/A18066). This work was supported by the Wellcome Trust (PG, 109967/Z/15/Z), the American Society of Hematology (PG, Global Research Award) and Cancer Research UK (PG, Advanced Clinician Scientist fellowship, C57799/A27964). K.R-P. was supported by the Academy of Medical Sciences (SBF004\1099) J.H.M.P. was supported by a research grant from Science Foundation Ireland (SFI) under Grant Number 16/RC/3948 and co-funded under the European Regional Development Fund and by FutureNeuro industry partners.

## Author Contributions

P.G. and K.W. designed the experiments. K.W. and L.S.D. performed laboratory assays. K.M.R. G.V.H. designed and performed the palmitate labelling assay. A.M.S.M., C.P., V.D. and S.C.J. helped with experiments. L.N.L. and P.G. performed computational analysis of published datasets. P.M. performed RNA sequencing analysis. K.W., G.G., C.M., K.R.K. and R.A. designed, performed or helped in vivo studies. J.H.M.P. provided reagents and input for UPR analysis. C.P. and K.R-P. provided input on methodology, reagents, experimental design and analysis of UPR. K.T. and G.S.V. designed and performed original CRISPR screen. K.W. and P.G. wrote and edited the manuscript. P.G. designed the study and supervised the project with B.J.P.H. mentorship/support at the start. All authors critically reviewed the manuscript and approved the final version.

## Disclosures of Conflicts of Interest

No conflicts to disclose

## Notes

### Competing Interest Statement

The authors have declared no competing interest.

### Summary of Updates

Various figures have been updated to reflect reviewer feedback. In particular, multiple experiments have been repeated in addition cell lines, the role of the ATF6 arm of the unfolded protein response has been strengthened and the role of ferroptosis as the form of cell death seen has been reinforced.

