## Supplemental Data and materials and methods for "Mannose metabolism inhibition sensitizes acute myeloid leukemia cells to cytarabine and FLT3 inhibitor therapy by modulating fatty acid metabolism to drive ferroptotic cell death"

Extended data

Including:

Materials and Methods

Extended data figures 1-8

### METHODS

**Cell culture**
MOLM13, MV411 and THP-1 cells were grown in 1640-RPMI (Gibco, 21875034) supplemented with 10% FBS (Sigma, F4135) and 1% penicillin/streptomycin (Gibco, 15140122) at 37°C with 5% $CO_2$, unless otherwise stated. For SEAHORSE analysis, media comprised of SEAHORSE RPMI (Agilent), without bicarbonate and FBS, supplemented with 11mM glucose and 2mM L-Glutamine (SEAHORSE media), unless otherwise stated. HEK-293T Phoenix cells were grown in DMEM (Gibco, 41966029) supplemented with 10% FBS (Sigma, F4135) and 1% penicillin/streptomycin (Gibco, 15140122) at 37°C with 5% $CO_2$ unless otherwise stated. For experiments where delipidated FBS was used, this was done by the adding of 2g of fumed silica to 100ml of heat inactivated FBS (Sigma, F4135) and stirring overnight at room temperature. The mixture was then centrifuged at 5000rpm for 25 minutes to pellet the silica and the supernatant retained. The supernatant was then filtered through a 0.2µm filter to ensure sterility before storage at -20°C for further use. Delipidated FBS was used in RPMI at a final concentration of 10%. Cells were only kept in delipidated FBS for 3 days as any longer culture of cells in these conditions lead to large scale cell death in all conditions.

**Lentiviral transduction**
293T-Pheonix cells were grown in DMEM (Gibco) containing 10% FBS and transfected with construct plasmid (NTshRNA and MPI shRNA1 and 2 TRIPZ inducible shRNA, Horizon Discovery, ATF6-YFP reporter fig. S5f), packaging construct (psPAX) and glycoprotein (pMD2G) using lipofectamine (Gibco, 18324010) for transient transfection. After 24 hours, medium was replaced by RPMI with 5% FBS before overnight incubation at 32°C with 5% $CO_2$. Media containing virus was harvested and filtered through a 0.45 µm filter to remove any cells 24 and 48 hours post media change and either used immediately or stored at -80°C in 1ml aliquots for further use.
For transduction of cell lines, 1 million cells were seeded in 1ml RMPI containing virus with 4µg/ml polybrene (Santa Cruz Biotechnology, TR1003) and spun for 90 minutes at 2500 rpm on 2 consecutive days. Cells were then placed in an incubator at 32°C for 4 hours before the addition of a further 3ml of RPMI with 5% FBS. Cells were then placed in an incubator at 37°C for 24 hours before the virus was removed by washing three times with sterile PBS.
Primary samples were cultured in StemSpan II (STEMCELLTechnologies, 09605) supplemented with 150 ng/ml SCF (Biolegend, 573908), 150 ng/ml Flt-3 (Biolegend, 550602), 10 ng/ml IL-6 (Biolegend, 570802), 25 ng/ml G-CSF (Biolegend, 578602), 20 ng/ml TPO (Biolegend, 763702), 1% HEPES (Gibco, 15630056) and 1% Penicillin/Streptomycin. All samples were cultured at 37°C with 5% $CO_2$. Cells were not spun and instead viral supernatant was added to a final volume of 10% of total culture volume. Samples were then cultured overnight before repeating. Virus was removed by washing three times with sterile PBS before continuing with culture as detailed below.
For MPI and NTshRNA transduced samples, cells were cultured in normal media with the addition of puromycin (1µg/ml final concentration, ThermoFisher, A1113803) for selection of transduced cells and doxycycline (1µg/ml final concentration, Sigma, D3072) for activation of the shRNA construct. Activation of the construct and transduction efficiency was determined by the percentage of RFP+ cells. Cells were then cultured as normal with the addition of 1µg/ml doxycycline.
For the ATF6 reporter, transduced cells were selected by treating cells with 1µM tunicamycin (Tocris, 3516) to induce YFP expression in transduced cells before cell sorting for the YFP positive population. Cells were then cultured as normal.

**Human samples**

The use of human samples, collection and publication of individual-level clinical data was approved by the East of England - Cambridge Research Ethics Committee REC 21/EE/0123. The conduct of the study was fully compliant with the ethical approval. Bone marrow and peripheral blood samples containing >80% blasts were collected from adult patients with AML at diagnosis or relapse and cryopreserved after mononuclear cell (MNCs) isolation using Lymphoprep™ (07851, STEMCELL Technologies) based density gradient centrifugation. Normal $CD34^+$ enriched samples were taken from myeloma patients at the time of peripheral blood stem cell collections. Samples were thawed and transduced via lentivirus with NTshRNA or MPIshRNA as described above.

After transduction was confirmed by the presence of RFP expression by flow cytometry, primary samples were cultured on a stroma of MS-5 mouse cells that had been irradiated in H5100 media (STEMCELL Technologies) supplemented with penicillin/streptomycin, interleukin-3, G-CSF and TPO at 20ng/ml final. For all assays, AML cells were removed from the co-culture and cultured without stroma. Assays were performed as described in viability section, however less cells were used in each case as numbers of cells in the primary samples was limited. To ensure that only transduced primary cells were removed and used for assays, RFP-positive percentage was assessed to be above 95%.

**Generation of CRISPR knockout clones**

The knock-out of genes was accomplished using the CRISPR/Cas9 system. For this, MOLM13, MV411 and THP-1 cell lines were transduced with Cas9 lentivirus (Addgene, lentiCas9-Blast, #52962), generating Cas9 expressing cell lines. Functional gRNA sequences were obtained from genome-wide gRNA library and ligated in the backbone, conjugating MPI gRNA with mCherry (Addgene, pKLV2-U6gRNA5(Bbsl)-PGKpuro2AmCherry-W, #67977). Virus for Cas9, MPI gRNA and NT gRNA were produced and transduced as described above. Obtaining single cell colonies was performed using methylcellulose media (H4531; STEMCELL Technologies, Cambridge, UK), where cells were cultured in semi-liquid conditions to minimize movement. Seeding cells in methylcellulose in a low density (~500 cells/mL) permitted separate colonies to form from a single cell. After picking the colonies from the assay and culturing them in small volumes, the generation of cell lines from a single parent all with the same construct was realized.

**CRISPR KO gRNA sequences**

NTgRNA - ATTTTCGTACCCTGGGACGC

MPIgRNA2 - AGGATCTTGGCATCCCCTC

MPIgRNA5 – CAATCAGGAACTGAAACTC

**SiRNA transfection**

ATF6 (Ambion, AM16708) and non-silencing control (Ambion, 4390843) siRNAs were resuspended in nuclease free water and stored at -20°C until further use. Molm13 cells were replated at a density of 200,000 cells/ml in normal RPMI with 10% FBS and 1% pen/strep 48 hours before transfection. On day of transfection cells were pelleted, washed twice with PBS and finally resuspended in 100µl buffer SF (Lonza, V4XC-2012). Cells were transferred to Lonza Nucelocuvettes and placed into the Lonza Nucelofector 4D X unit and transfected using program DJ-100. Cells were then removed and added to 2ml of pre-warmed RPMI with 10% FBS and 1% pen/strep. Cells were taken for RNA extraction and qPCR analysis after 48 hours and plated for annexin analysis at the same time.

**Treatments**

AC220 (Insight Biotechnology, HY13001) was dissolved in DMSO and cells were treated at 1nM final concentration, unless otherwise stated. Mannose (Sigma, 112585) was dissolved in sterile water and cells were treated at 100µM final concentration, unless otherwise stated. Etomoxir (Cayman Chemicals, 11969) was dissolved in DMSO and cells were treated at 10µM final concentration for all apoptosis and lipid peroxidation assays and 50µM for all SEAHORSE experiments, unless otherwise stated. MLS0315771 (Insight Biotechnology, HY112945) was dissolved in DMSO and cells were treated at a final concentration of 1µM, unless otherwise stated. Ceapin A7 (Sigma, SML2330) was dissolved in DMSO and cells were treated at a final concentration of 1µM, unless otherwise stated. Fenofibrate (Sigma, F6020) was dissolved in DMSO and cells were treated at a final concentration at 10µM, unless otherwise stated. Ferrostatin-1 (Sigma, SML0583) was dissolved in DMSO and cells were treated at a final concentration of 5µM, unless otherwise stated. Erastin (Sigma, E7781) was dissolved in DMSO and cells were treated at a final concentration of 4µM, unless otherwise stated. Sulfosuccinimidyl Oleate (SSO, Cayman Chemicals, 11211) was dissolved in DMSO and cells treated at a final concentration of 100µM. Arachidonic acid (Sigma, 10931), palmitic acid (Acros organics, 129702500) and oleate (Sigma, O7501) were dissolved in isopropanol, with cells treated in concentrations as outlined below. AA147 (Tocris, 6759) was dissolved in DMSO and cells were treated at a final concentration of 5µM. Tunicamycin (Tocris, 3516) was dissolved in DMSO and cells were treated at a final concentration of 1µM. ZVAD (Sigma, V116) was dissolved in DMSO and cells were treated at a final concentration of 20µM. Necrostatin-1 (Sigma, N9037) was dissolved in DMSO and cells were treated at a final concentration of 20µM. GSK2656157 (SelleckChem, S7033) was dissolved in DMSO and cells were treated at a final concentration of 200nM. MKC-3946 (SelleckChem, S8286) was dissolved in DMSO and cells were treated at a final concentration of 1µM. Dimethyl alpha-ketoglutarate (Sigma, 349631) was used at a final concentration of 4mM. RSL3 (SelleckChem, S8155) was dissolved in DMS and used at a final concentration of 100nM unless otherwise stated.

**Viability Assay**

Around 1 million cells per condition were harvested by centrifugation at 500xg, before washing twice with PBS at 4°C. Cells were then resuspended in annexin V binding buffer (BioWorld, 21720002-1) and annexin V-FITC (BioLegend, 640945) and 7-AAD (BD Biosciences, 559925) were added. Positive control cells for single colour compensation of the FITC and 7-AAD were generated by heating cell suspensions for 3 minutes at 70°C. Cells were incubated for at least 15 minutes at 4°C, protected from light. For analysis of live cells by DAPI exclusion cells were harvested as described above and resuspended in PBS. DAPI at a final concentration of 0.5µg/ml was added. Cells were then kept at 4°C protected from light until analysis. Cells were then analysed using LSR Fortessa (BD Biosciences) instruments. Downstream analysis was performed using FlowJo version 10.6.1 (BD Biosciences).

**Cell counting**

Cells were counted either using a CASY cell counter (Cambridge Biosciences) or a LUNA automated cell counter (Logos Biosystems). For LUNA cell counting, cell samples were mixed in a 1:1 ratio with 0.4% trypan blue solution.

**Competition assays**

125000 Cas9/KO-mCherry (NTgRNA or MPIgRNA) per ml were added along with 125000 Cas9 cells with no mCherry (parental) per ml and were cultured in the presence or absence of AC220 (1nM). At days 3, 6 and 9 the ratio between mCherry positive and negative cells were measured by flow cytometry using LSR Fortessa (BD Biosciences) instruments, with data normalised to the NT:parental ratio at day 3. Downstream analysis was performed using FlowJo version 10.6.1 (BD Biosciences).

**IC50 calculations**

50000 NT gRNA, MPI gRNA2 and MPI gRNA5 MOLM13 cells were seeded and treated with increasing concentrations of AC220 in triplicate. After 3 days cell proliferation was measured using CellTiter 96 AQueous Non-Radioactive Cell Proliferation Assay (Promega, G5421) and normalised to cells treated with vehicle only as 100% proliferation.

**SEAHORSE MitoStress test**

The SEAHORSE MitoStress test was performed essentially per manufacturer's instructions. In short, cells were grown and treated as described above. The SEAHORSE cartridge was hydrated with sterile water overnight in a 37°C incubator with no $CO_2$, before the water being removed and replaced with SEAHORSE calibrant solution (Agilent, 100840-000) for at least 1 hour before the cartridge is put into a SEAHORSE XF instrument. The cell plate was coated with Corning CellTak (Fisher, can no CB40240) adhesion solution at 22.4µg/ml concentration in sodium bicarbonate at pH 8 for 15 minutes at room temperature before washing twice with sterile water. $10^5$ cells were harvested and washed with PBS before resuspension in 50µl of pre-warmed SEAHORSE RPMI media (Agilent, 103576-100) and adding onto the cell plate. Cells were adhered by centrifugation at 100xg for 1 minute without braking, before incubation at 37°C for 15 minutes in an incubator with no $CO_2$. 130µl of additional SEAHORSE media along with etomoxir at a final concentration of 50µM (only to etomoxir treatment wells) was added to wells and incubated for a further 15 minutes. The drugs were added to the cartridge in the following manner: port A – oligomycin, 1µM final concentration, port B – FCCP, 0.3µM final concentration, port C – rotenone/antimycin A, 0.5µM final concentration (Agilent, 103015-100), all of which were prepared in SEAHORSE media. The assay was set up on the SEAHORSE XF machine with a 2 minute mixing, 2 minute wait and 2 minute measuring for each well at each timepoint.

For palmitate SEAHORSE assays it was performed essentially as above, but cells were cultured overnight in RPMI media lacking FBS and glutamine, supplemented with 50µM palmitate-BSA or BSA (Sigma, A8806) only. SEAHORSE media was without glutamine and palmitate (50µM) was added to the + palmitate conditions. The rest of the assay was run as described above.

**Bodipy uptake assay**

C1-Bodipy C12 500/510 (Thermo Fisher, D3823) was added to cells in normal culture conditions at a final concentration of 1µM, concurrent with other treatments before being placed in an incubator at 37°C and 5% $CO_2$ for 24 hours. Cells were then pelleted and washed twice with PBS before resuspension in PBS and flow cytometry analysis using LSR Fortessa (BD Biosciences) instruments. Downstream analysis was performed using FlowJo version 10.6.1 (BD Biosciences).

**Bodipy Lipid peroxidation assay**

1 million cells of each condition were harvested and the positive control was treated with 100µM tetrabutyl hydroperoxide for the duration of the Bodipy labelling. Bodipy 581/591 lipid peroxidation sensor (Thermos Fisher, D3861) was added to a final concentration of 4µM and cells were incubated for 45 minutes at 37°C with gentle shaking (300rpm), protected from light. Cells were then removed, washed twice with PBS and analysed by flow cytometry using LSR Fortessa (BD Biosciences) instruments. Downstream analysis was performed using FlowJo version 10.6.1 (BD Biosciences).

**Bodipy neutral lipid stain**

Cells were grown as normal with the addition of fatty acids-BSA or BSA alone as vehicle at the following concentrations: palmitic acid: 50µM, oleate: 50µM and arachidonic acid 1µM for 24 hours. Around 1 million cells of each condition were then harvested and washed twice in PBS, before resuspension in PBS and incubating the samples with 2.5µg/ml Bodipy 493/503 (ThermoFisher, D3922) for 20 minutes at 37°C with agitation. Samples were then washed twice with PBS and resuspended in PBS and then kept on ice before analysis by flow cytometry using LSR Fortessa (BD Biosciences) instruments. For data interpretation, cells of all conditions were grown in normal media with BSA only. These cells were harvested and stained as above and the mean fluorescence intensity (MFI) was measured and subtracted from lipid supplemented conditions as a baseline of intracellular lipid content. Downstream analysis was performed using FlowJo version 10.6.1 (BD Biosciences).

**Metabolomic analysis**

Cells were treated with AC220 (1nM) and mannose (100µM) for 48 hours before 30 million cells were harvested and washed. Pellets were snap frozen and sent to Metabolon (Morrisville, NC) for LC-MS analysis. After extraction of metabolites, samples were split into four aliquots to be analysed in acidic positive, acidic negative and basic negative eluted from a UPLC C18 column (Waters) with the fourth aliquot separated with negative ionisation on a UPLC HILIC column (Waters). All analysis was performed on a Q-Exactive mass spectrometer (Thermo Scientific) coupled to an ACQUITY UPLC system (Waters) coupled to a HESI-II electrospray ionisation source. MS operated at 35000 mass resolution with scan range between 70 and 1000 m/z. All extraction and analysis of RAW data was performed in house by Metabolon. Raw counts were normalised by Bradford protein concentration and then each biochemical is rescaled to the median levels of each biochemical across all 6 conditions which is set as reference. *Z* scores are then calculated as the difference between the observed value in one condition minus the mean, all divided by the standard deviation, i.e. $x$-µ/σ. Targeted metabolomic analysis following U-$^{13}C_6$-glucose labelling was performed as described previously[9].

**Palmitate labelling**

Cells were cultured in Plasmax media (Ximbio, 156371) supplemented with 5% dialysed FBS (Sigma). U-$^{13}C_{16}$-Palmitic acid (CK Isotopes, CLM409) was dissolved in isopropanol and added to cell culture concurrent with treatments at a final concentration of 50µM for 24 hours. 500,000 cells were harvested washed twice with ice cold PBS before extraction in solvent containing 50% methanol (Fisher, 10675112), 30% acetonitrile (Fisher, 10407440) and 20% $H_2O$ (All HPLC grade, Fisher, 10777404). Samples were then centrifuged and the supernatant retained. Samples were kept at -80°C until analysis.

For liquid chromatography a ZIC-pHILIC guard column (SeQuant, 20 × 2.1 mm) and ZIC-pHILIC column (SeQuant, 150 × 2.1 mm, 5 µm, Merck KGaA) was used on a Thermo Scientific UltiMate 3000 HPLC system. The aqueous mobile-phase solvent consisted of a 0.1% ammonium hydroxide -20 mM ammonium carbonate solution and and the organic mobile phase was acetonitrile. A linear biphasic LC gradient was executed, staring from 80% organic phase to 80% aqueous phase. The gradient took 15 minutes for a total run time of 22 minutes. The flow rate was set to 200 µl/min with column temperature being maintained at 45°C. The mass spectrometer used was a qExactive Orbitrap Mass Spectrometer (Thermo Fisher Scientific). The MS operated in polarity switching mode. Pre-analysis calibration was carried out for both ionization mode using a custom CALMIX and a low m/z range tune file was used. Full scan (MS1) data was acquired during polarity switch in profile mode. The resolution was 35,000 (at m/z range 75–1000). Automatic gain control was set to target of 1x106 with a max fill time of 250 ms. The parameters for spray voltages are as follows: +4.5 kV (capillary +50 V, tube: +70 kV, skimmer: +20 V) and −3.5 kV (capillary -50 V, tube: -70 kV, skimmer: -20 V). The s-lens RF level was set to 50 for the front optics. The probe temperature was set at 50̊ C and capillary temperature set at 375̊ C The sheath gas flow rate was 25 and auxiliary gas flow rate was 15. Sweep gas flow was set to rate of 1. The mass accuracy was sub 5 ppm and data was acquired using Thermo Xcalibur 4.3.73.11 software.

The peak areas of each metabolite and their respective isotopologues were quantified using Thermo Tracefinder 4.1. Metabolites were identified by accurate mass of the singly charged ion and by retention times of authentic standards on the pHILIC column. The commercially standard compound mix (Merck: MSMLS-1EA) had been analysed previously on our LCMS system to determine accurate ion masses and retention times. Data were processed and corrected for natural abundance through Autoplotter. (Pietzke and Vazquez, 2020, Cancer Metab. doi: 10.1186/s40170-020-00220-x.).

**Immunocytochemistry**

Approximately 1 million cells for each condition were harvested and washed twice with PBS before fixing with 4% paraformaldehyde in PBS for 10 minutes at room temperature. Cells were then washed before permeabilisation with 0.3% triton x-100 in PBS for 15 minutes at room temperature, before a PBS wash and blocking with 0.2% fish skin gelatin for 1 hour at room temperature with agitation. Cells were incubated with primary antibody diluted in blocking solution (ATF6, 1:200, Santa Cruz biotech, SC166659) for 1 hour at room temperature with agitation before 5 washes with PBS. Samples were then incubated with secondary antibody (AlexFluor 488, goat anti-rabbit, 1:1000, Thermo, A32731) protected from light before 5 washes with PBS. Glass slides were coated with CellTak solution (22.4µg/ml) at room temperature for 15 minutes before washing the slides twice with PBS. Cells were then pipetted onto the CellTak coated slides and left for 30 minutes at room temperature before the addition of DAPI Fluoromount (Southern Biotech, 0100-20) solution and covering with coverslips. Slides were then kept at 4°C until imaging. All imaging was done on a Zeiss LSM 710 scanning confocal microscope at 40x or 63x magnification. Further image analysis and processing was performed using ImageJ, including use of the Coloc2 plugin for colocalisation analysis. For 4-HNE staining, the procedure was performed essentially as above, with anti-4-HNE antiserum (Alpha Diagnostic International, HNE11-S) as the primary antibody. Further analysis was performed with the FIJI software package for ImageJ, using the Coloc2 plugin for colocalisation. For enhancing contrast, saturated pixels were set to 0.3 or 0.5% and was performed on the whole image at once. For images where more than 0.3 or 0.5% of pixels were already saturated, this contrast enhancement had no visible effect.

**Staining of surface proteins for flow cytometry**

Approximately 1 million cells for each condition were harvested and washed twice with PBS before resuspension in PBS. FC block (Biolegend, 422301) was then added for 20 minutes at room temperature before the addition of primary antibody at a 1:100 dilution before incubation at 4°C for 30 minutes. Cells were then washed 3 times with PBS before resuspension in PBS. Samples were then stored either on ice or at 4°C protected from light until analysis by flow cytometry using LSR Fortessa (BD Biosciences) instruments.

The following antibodies were used: anti-human CD45-FITC (BioLegend, 304005), anti-mouse CD45-APC (BioLegend, 157605), anti-SLC7a11 (SantaCruz Biotech, SC98552). Downstream analysis was performed using FlowJo version 10.6.1 (BD Biosciences).

**Lectin staining of surface glycoproteins**

Approximately 1 million cells for each condition were harvested and washed twice with PBS before resuspension in PBS. Lectin from *Lycopersicon esculentum* (tomato) FITC conjugate (Sigma, L0401) was added in a 1:100 dilution before incubation for 1 hour at 4°C, protected from light. Cells were then washed twice with PBS, resuspended in PBS before flow cytometry analysis using LSR Fortessa (BD Biosciences) instruments. Downstream analysis was performed using FlowJo version 10.6.1 (BD Biosciences).

**Proteostat staining**

Approximately 1 million cells per condition were collected and washed twice with PBS before fixation for 10 minutes at room temperature by resuspension in 4% paraformaldehyde. Cells were then washed twice with PBS and permeabilised with 0.3% triton X-100 in PBS for 15 minutes before a further wash and resuspension in assay buffer from the proteostat protein aggregation kit (Enzo life sciences, ENZ51023) with proteostat stain (1:5000). Cells were then kept on ice before analysis.

**Western blot analysis**

$10^6$ Cells were lysed in Cell Signalling technololgies lysis buffer (Cell Signalling) supplemented with protease and phosphatase inhibitors. DNA was sheared by passing the sample through a fine gauge needle 8 times before centrifugation for 10 minutes at 14000xg. The supernatants were retained and stored at -20°C before further use. NuPAGE sample buffer (Invitrogen) was added to samples before separation on 4-12% polyacrylamide gels (Invitrogen, NP0321). Proteins were then transferred to PVDF membranes followed by blocking with 5% milk/Tris Buffered Saline with 0.1% tween (TBS-T), incubation with primary antibodies in 5% BSA/TBS-T overnight at 4°C then corresponding secondary antibodies in 5% milk/TBS-T for 1 hour at room temperature before incubation of the membranes with ECL (BioRad, . 1705061) and visualisation using an Amersham Imager (GE Healthcare). Further analysis was performed in ImageJ. Primary antibodies were used at the following concentrations: ATF6 (Abcam, AB122897), 1:500, total Histone 3 (Cell Signalling, 44995), 1:5000, MPI (Santa Cruz Biotech, SC393477), 1:1000, PARP (Cell Signalling, 56255), 1:500, EIF2α (Cell Signalling, 9722), 1:1000, phosphor-EIF2α (Ser51, Cell Signalling, 9721), 1:1000.

**Quantitative RT-PCR (qPCR)**

RNA was extracted from ~1 million cells using the RNeasy mini kit (Qiagen, 74106) as per the manufacturer's instructions. RNA concentration was then measured using a nanodrop and immediately reverse transcribed using the High-Capacity cDNA Reverse Transcription Kit (ThermoFisher, . 4368814) as per the manufacturers intstructions. cDNA was then stored at -20°C until use. For qPCR 20ng of cDNA was added to 5ul of PowerUp SYBR Green MasterMix (ThermoFisher, A25742) with 250nM of forward and reverse primers for the gene of interest. Samples in a 384 well plate were then analysed by BioRad CFX384 Real-Time System (BioRad) with the following protocol: 95°C for 2 minutes, 95°C for 5 seconds, 60°C for 30 seconds when the fluorescence was measured before returning to step 2 and repeating 39 times, 95°C for 5 seconds, 65°C for 5 seconds, fluorescence measurement, 95°C for 30 seconds and holding at 16°C forever.

**qPCR primer sequences**

CPT1A - Forward: CAAACTGGACCGGGAGGAAA; Reverse: TGTGCTGGATGGTGTCTGTC.

PPARA - Forward: AGCTGTCACCACAGTAGCTT; Reverse: GGAACTCTTCAGATAACGGGCT.

ACOX1 - Forward: CGCCGAGAGATCGAGAACAT; Reverse: GCACTTTTCCTGACAGCCAC

MPI - Forward: CTGCCGGGAAAGGCATACG; Reverse: AGCAATCCACTGGCTTAGGG

ERO1B - Forward: AGAGAACTGTTTCAAGCCTCG; Reverse: TCCAGACACAAACCTTCTAGCC

HERPUD1 - Forward: CGAGATTGGTTGGATTGGACC; Reverse: CACCCAACGTGATGCAGGTA

XBP1 spliced – Forward: TTGCTGAAGAGGAGGCGGAA; Reverse: CTGCACCTGCTGCGGACTCAG

XBP1 total – Forward: TTCCGGAGCTGGGTATCTCA; Reverse: GAAAGGGAACCCCCGTATCC

ATF6 – Forward: ATGAAGTTGTGTCAGAGAACC; Reverse: CTCTTTAGCAGAAAATCCTAG

**RNA sequencing**

Cells were treated with AC220 (1nM) and mannose (100µM) for 24 or 72 hours as indicated. RNA was extracted and purified with the RNAeasy kit (Qiagen, 74106) as per manufacturer's instructions. Concentration was measured by NanoDrop (ThermoFisher). Ribosomal depletion was performed using the QIAseq FastSelect™ RNA Removal Kit - 24samples; HUMAN (Qiagen: THS-001Z-24). Library preparation was performed with the dUTP directional kit NEBNext® Ultra8482 II Directional RNA Library Prep Kit for Illumina®. Samples were analysed on a NovaSeq6000 S1 flowcell as a paired end

150bp (PE150) run. We followed recommended guidelines in the analysis of RNA-seq data for quality control, read mapping, quantification of gene expression, assessment of reproducibility among biological replicates, and differential gene expression (Conesa et al., 2016, Genome Biology, doi: 10.1186/s13059-016-0881-8.). First, TrimGalore 0.4.5 was used with default parameters and paired-end mode for quality-trimming and filtering of read sequences. Then, STAR 2.6.1 (Dobin et al, 2013, Bioinformatics, doi: 10.1093/bioinformatics/bts635) was used to align the reads to the reference human genome assembly GRCh38 using the Ensembl release 95 annotation as reference transcriptome. Proportion of uniquely mapped reads was > 91% in all samples. FeatureCounts 1.6.3 was run on paired-end reads to count fragments in annotated gene features, with parameters '-p -T 4 -t exon -g gene_id' (Liao et al., 2014). The R/Bioconductor package DESeq2 was then used to perform differential gene expression analyses between samples and conditions (Love et al., 2014, Genome Biology, doi: 10.1186/s13059-014-0550-8). Raw data is available at https://www.ebi.ac.uk/arrayexpress/experiments/E-MTAB-11750.

**Gene-set Enrichment Analysis (GSEA) analysis**

RNA-sequencing data was analysed for GSEA using the Broad Institute software (http://software.broadinstitute.org/gsea/index.jsp). For GSEA analysis normalised read counts for all conditions were used and genes were ranked using the signal-to-noise metric and FDR and NES were calculated using 1000 gene-set permutation.

**Public dataset analysis**

Public gene array datasets GSE6891, E-TABM 1029, GSE12417, GSE15434, GSE13159, GSE10358, GSE37642, GSE76009, GSE30377, GSE83533 and RNA-Seq from AML TCGA (data obtained from https://www.cbioportal.org/), BeatAML dataset (data obtained http://www.vizome.org/) and from manuscript 10.1056/NEJMoa1808777 (NEJM WUSM) were used to perform various analysis. For MPI expression, normalised expression was used and compared between different genotypes ($FLT3^{ITD}$ vs $FLT3^{WT}$) or patient samples (diagnosis vs relapse). For analysis of MPI level based on AML karyotype pre normalised data from GEO GSE147515 was plotted based on AML karyotype and MPI level. For GSEA analysis, within relevant datasets, the expression profile patients expressing high (i.e. above median) levels of MPI was compared to that of patients expressing low (i.e. below median) MPI levels. Genes were ranked using the signal-to-noise metric and FDR and NES were calculated using 1000 phenotype permutation.

***In vivo* experiments**

$3x10^6$ luciferase expressing MOLM13 cells transduced with non-targeting CRISPR guide (NT-gRNA) or MPI KO CRISPR guide (MPI-gRNA) were transplanted into male 8-12 week old NSG (NOD.Cg-*Prkdc*$^{scid}$ *Il2rg*$^{tm1Wjl}$/SzJ) mice which were sublethally irradiated (2 Gy). Mice were treated by oral gavage with AC220 (5mg/kg) or vehicle (22% hydroxypropyl-β-cyclodextrin/0.3% DMSO) for 5 days. Survival time was measured from time of transplantation until time mice had to be culled due to overt clinical symptoms. IVIS bioluminescence imaging (PerkinElmer) was used to confirm and track disease dissemination. In brief, D-luciferin (Cat#122799, Perkin Elmer) was administered by intraperitoneal (IP) injection as per the manufacturer's recommendation, followed by IVIS bioluminescence imaging under anesthesia (Isoflurane). Bioluminescence as a surrogate for tumour burden, was quantified using Living Image Software (version 4.7.2, PerkinElmer).

8 week old male NBSGW (NOD.Cg-*Kit*$^{W-41J}$ *Tyr* $^{+}$ *Prkdc*$^{scid}$ *Il2rg*$^{tm1Wjl}$/ThomJ) mice had either 5000 THP-1 NTgRNA or THP-1 MPIgRNA5 cells infused by intravenous injection via the tail vein. 1 week after this AML engraftment was measured by flow cytometry based on percentage of human CD45+ cells in the peripheral blood. The next day mice were treated with 50mg/mg cytarabine (Sigma, 251010) in PBS

or PBS alone (vehicle) for 7 days via intraperitoneal injection. Survival time was measured from time of transplantation until time mice had to be culled due to overt clinical symptoms.

Bone marrow tissue was harvested by dissection of the leg bones of the mouse before the bones were flushed with PBS to obtain the cells. After collection 2ml of red blood cell lysis buffer (ThermoFisher, 00430054) was added and incubated for 5 minutes at room temperature before inactivation by the addition of 10 volumes of PBS. Cells were then pelleted for further flow cytometry analysis as outlined previously above.

10 week old male NBSGW (NOD.Cg-*Kit*$^{W\text{-}41J}$ *Tyr* $^{+}$ *Prkdc*$^{scid}$ *Il2rg*$^{tm1Wjl}$/ThomJ) mice had either 5000 THP-1 NTgRNA or THP-1 MPIgRNA5 cells infused by intravenous injection via the tail vein. 1 week after this AML engraftment was measured by flow cytometry based on percentage of human CD45+ cells in the peripheral blood. The next day mice were treated with 50mg/mg cytarabine (Sigma, 251010) in PBS for 7 days with or without liproxstatin (10mg/kg) for 10 days via intraperitoneal injection. Survival time was measured from time of transplantation until time mice had to be culled due to overt clinical symptoms. For 4-HNE assessment a cohort of 3 mice in each treatment group was culled soon after treatment and bone marrow harvested. Thereafter human CD45+ leukaemic cells were sorted for immunocytochemistry as described above.

**Statistical analysis**

All data was visualised in Prism 8.4 (Graph Pad), with statistical tests outlined in figure legends for individual analyses. All data are shown as mean ± standard error of the mean, unless otherwise stated.

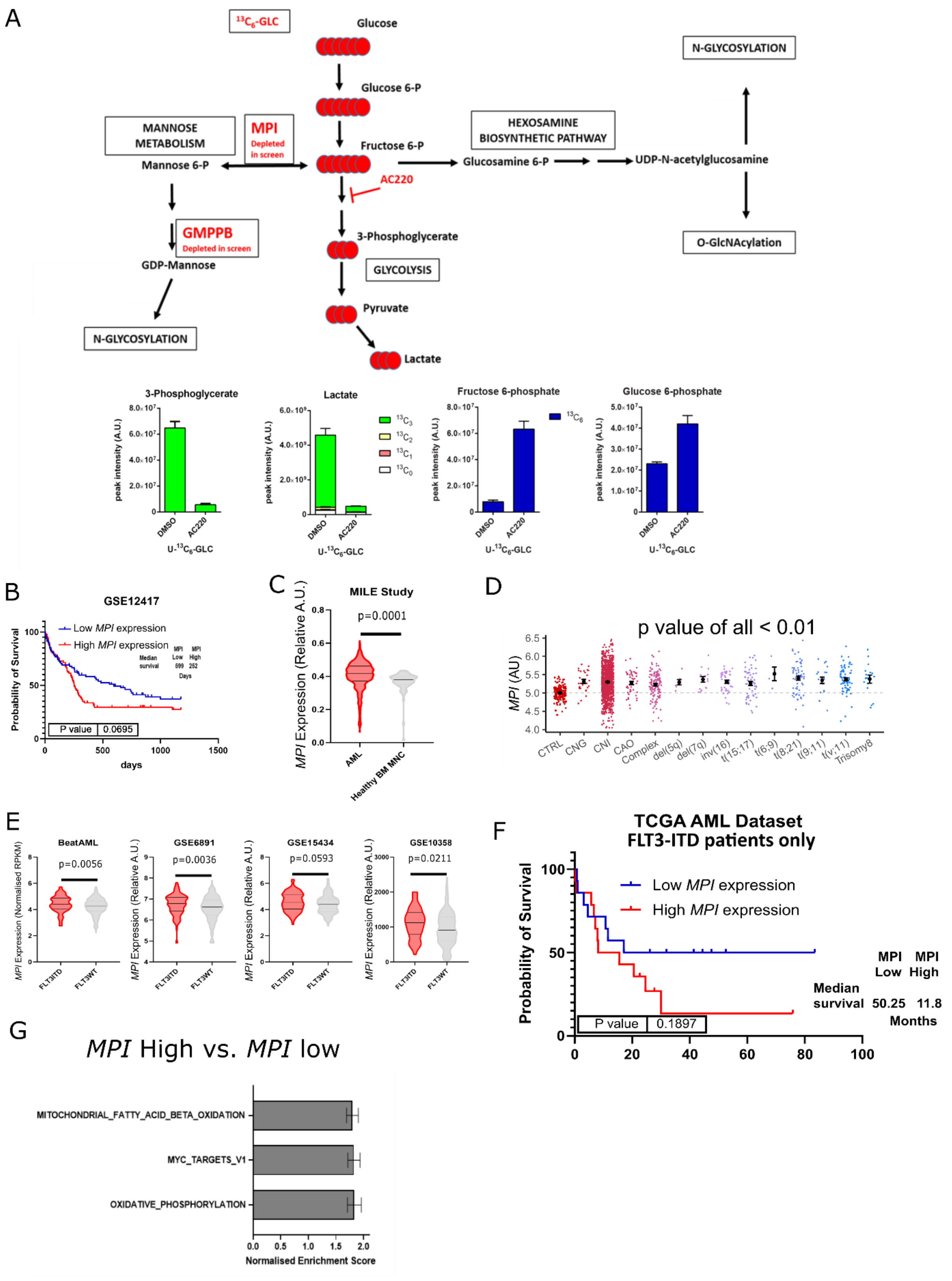
A
13C6-GLC
Glucose
Glucose 6-P
Fructose 6-P
MANNOSE METABOLISM
MPI Depleted in screen
Mannose 6-P
AC220
HEXOSAMINE BIOSYNTHETIC PATHWAY
Glucosamine 6-P
UDP-N-acetylglucosamine
N-GLYCOSYLATION
O-GlcNAcylation
GMPPB Depleted in screen
GDP-Mannose
N-GLYCOSYLATION
3-Phosphoglycerate
GLYCOLYSIS
Pyruvate
Lactate
3-Phosphoglycerate
Lactate
Fructose 6-phosphate
Glucose 6-phosphate
peak intensity (A.U.)
DMSO
AC220
U-13C6-GLC
B
GSE12417
Low MPI expression
High MPI expression
Probability of Survival
P value 0.0695
days
C
MILE Study
p=0.0001
MPI Expression (Relative A.U.)
AML
Healthy BM MNC
D
p value of all < 0.01
MPI (AU)
CTRL
CNG
CNI
CAO
Complex
del(5q)
del(7q)
inv(16)
t(15;17)
t(6;9)
t(8;21)
t(9;11)
t(v;11)
Trisomy8
E
BeatAML
p=0.0056
GSE6891
p=0.0036
GSE15434
p=0.0593
GSE10358
p=0.0211
MPI Expression (Normalised RPKM)
MPI Expression (Relative A.U.)
FLT3ITD
FLT3WT
F
TCGA AML Dataset
FLT3-ITD patients only
Low MPI expression
High MPI expression
Probability of Survival
P value 0.1897
MPI Low
MPI High
Median survival 50.25 11.8
Months
G
MPI High vs. MPI low
MITOCHONDRIAL_FATTY_ACID_BETA_OXIDATION
MYC_TARGETS_V1
OXIDATIVE_PHOSPHORYLATION
Normalised Enrichment Score

**Extended data Figure 1**
**MPI is more highly expressed in AML cells compared to normal bone marrow**

**A** – Schematic of glycolysis and mannose metabolism with targets of AC220 and location of MPI and the downstream enzyme GMPPB (also depleted in reference 9) highlighted (top). Levels of glycolysis intermediates in MV411 cells treated with 1nM AC220 or vehicle grown in media containing uniformly-labelled $^{13}$Carbon(U-$^{13}C_6$)-glucose(GLC) measured by LC-MS analysis (bottom – data produced in reference 9); **B** - Kaplan-Meier curves comparing survival of patient from the GSE12417 dataset separated by the top 50% and bottom 50% of *MPI* expression. Log-rank (Mantel-Cox) test; **C** – Violin plots of *MPI* expression (relative) in AML samples compared to healthy bone marrow mononuclear cell samples from MILE (GSE13204) study. Unpaired t-test; **D** – Levels of *MPI* RNA in various AML subtypes when compared to control from GSE147515. **E** - Violin plots of *MPI* expression (normalised or relative) in FLT3-ITD AML samples compared to FLT3 WT AML samples from Beat AML dataset (left), GSE6891 dataset (middle left), GSE15434 dataset (middle right) and GSE10358 dataset (right). Unpaired t-test; **F** – Kaplan-Meier curves comparing survival of patient from the FLT3-ITD patients in the TCGA dataset separated by the top 50% and bottom 50% of *MPI* expression. Log-rank (Mantel-Cox) test; **G** - 3 significantly enriched gene signatures in *MPI* high expressing samples from 7 AML datasets (GSE6891, E-TABM1029, GSE12417, GSE15434, TCGA, GSE10358 and GSE13159) combined.

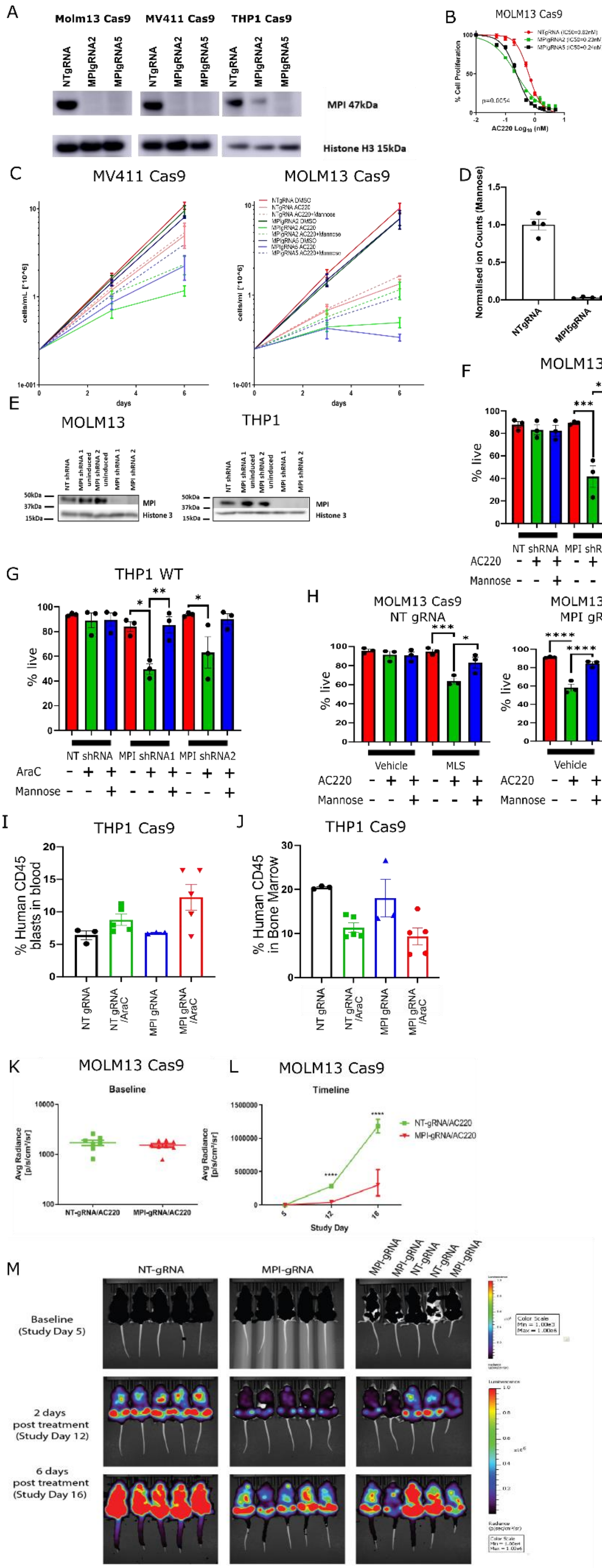
A
Molm13 Cas9
MV411 Cas9
THP1 Cas9
NTgRNA
MPIgRNA2
MPIgRNA5
MPI 47kDa
Histone H3 15kDa
B
MOLM13 Cas9
% Cell Proliferation
AC220 Log10 (nM)
C
MV411 Cas9
MOLM13 Cas9
cells/mL [*10^6]
days
D
Normalised ion Counts (Mannose)
NTgRNA
MPI5gRNA
E
MOLM13
THP1
NT shRNA
MPI shRNA 1
MPI shRNA 2
uninduced
MPI
Histone 3
50kDa
37kDa
15kDa
F
MOLM13 WT
% live
NT shRNA
MPI shRNA1
MPI shRNA2
AC220
Mannose
G
THP1 WT
% live
NT shRNA
MPI shRNA1
MPI shRNA2
AraC
Mannose
H
MOLM13 Cas9
NT gRNA
MOLM13 Cas9
MPI gRNA5
% live
Vehicle
MLS
AC220
Mannose
I
THP1 Cas9
% Human CD45 blasts in blood
NT gRNA
NT gRNA /AraC
MPI gRNA
MPI gRNA /AraC
J
THP1 Cas9
% Human CD45 in Bone Marrow
NT gRNA
NT gRNA /AraC
MPI gRNA
MPI gRNA /AraC
K
MOLM13 Cas9
Baseline
Avg Radiance [p/s/cm²/sr]
NT-gRNA/AC220
MPI-gRNA/AC220
L
MOLM13 Cas9
Timeline
NT-gRNA/AC220
MPI-gRNA/AC220
Study Day
M
NT-gRNA
MPI-gRNA
Baseline (Study Day 5)
2 days post treatment (Study Day 12)
6 days post treatment (Study Day 16)
Color Scale
Luminescence
Radiance

**Extended data figure 2**
**MPI CRISPR KO, shRNA KD and chemical inhibition show a consistent phenotype**

**A** – Western blot for MPI in MOLM13 (left), MV411 (middle) and THP1 (right) cells transduced with NT gRNA (left lane), MPI gRNA2 (middle lane) and MPI gRNA5 (right lane) showing MPI knockout with total Histone H3 as a loading control; **B** – IC50 dose curves for AC220 dosing of MOLM13 NT gRNA, MPI gRNA2 or MPI gRNA5. RM one-way ANOVA for treatment effect; **C** – Cell proliferation of MV411 (left) and MOLM13 (right) NT gRNA, MPI gRNA2 or MPI gRNA5 cells treated with vehicle, AC220 (1nM) or AC220 (1nM) and mannose (100µM) as indicated; **D** – Normalised mannose ion counts showing intracellular mannose levels of NTgRNA and MPIgRNA5 MOLM13 cells from untargeted metabolomics; **E** – Western blot for MPI in MOLM13 (left) and THP1 (right) cells transduced with NT shRNA (left lane), MPI shRNA1 uninduced (2$^{nd}$ lane), MPI shRNA2 uninduced (3$^{rd}$ lane), MPI shRNA1 induced (4$^{th}$ lane), MPI shRNA2 induced (5$^{th}$ lane) showing MPI knockdown with total Histone H3 as a loading control. Induction was carried out by treatment of cells with 1µg/ml doxycycline for 72 hours before cell harvest; **F** - Percentage of live cells of WT MOLM13 cells transduced with either NTshRNA, MPIshRNA1 or MPIshRNA2 as indicated treated with vehicle, AC220 (1nM), mannose (100µM) or in combinations as indicated. Treated for 6 days, N=3, 1 way Anova with Tukey's correction for multiple comparisons; **G** - Percentage of live cells of NT shRNA, MPI shRNA1 and MPI shRNA2 THP1 cells treated with vehicle, mannose (100µM), AraC (1µM) or AraC (1µM) and mannose (100µM), as indicated. Treated for 3 days, N=4 for MV411, N=3 for MOLM13, 1 way Anova with Tukey's correction for multiple comparisons; **H** - Percentage of live cells of MOLM13 NT gRNA (left) and MPI gRNA5 (right) cells treated with vehicle, MLS0315771 (1µM), AC220 (1nM), mannose (100µM) or in combinations as indicated. Treated for 6 days, N=3, 1 way Anova with Tukey's correction for multiple comparisons; **I** – Percentage of human CD45+ (THP1) cells in peripheral blood of NBSGW mice prior to the start of Cytarabine treatment; **J** – Percentage of human CD45+ (THP1) cells in bone marrow of NBSGW mice at the terminal timepoint after treatment with vehicle or cytarabine (50mg/kg for 7 days); **K** – Baseline engraftment of NT gRNA and MPI gRNA5 MOLM13 cells in NSG mice as measured by bioluminescence on day 5 post-transplant; **L** – Bioluminescence of NT gRNA and MPI gRNA5 MOLM13 cells engrafted into NSG mice over time after 5 days of treatment with AC220 (5mg/kg). Unpaired t-test; **M** – Visual representation of luminescence of NSG mice engrafted with NT gRNA or MPI gRNA5 MOLM13 cells at baseline and after treatment with AC220 (5mg/kg) over time. For all panels, ns = not significant, *=p<0.05, **=p<0.01, ***=p<0.005, ****=p<0.001.

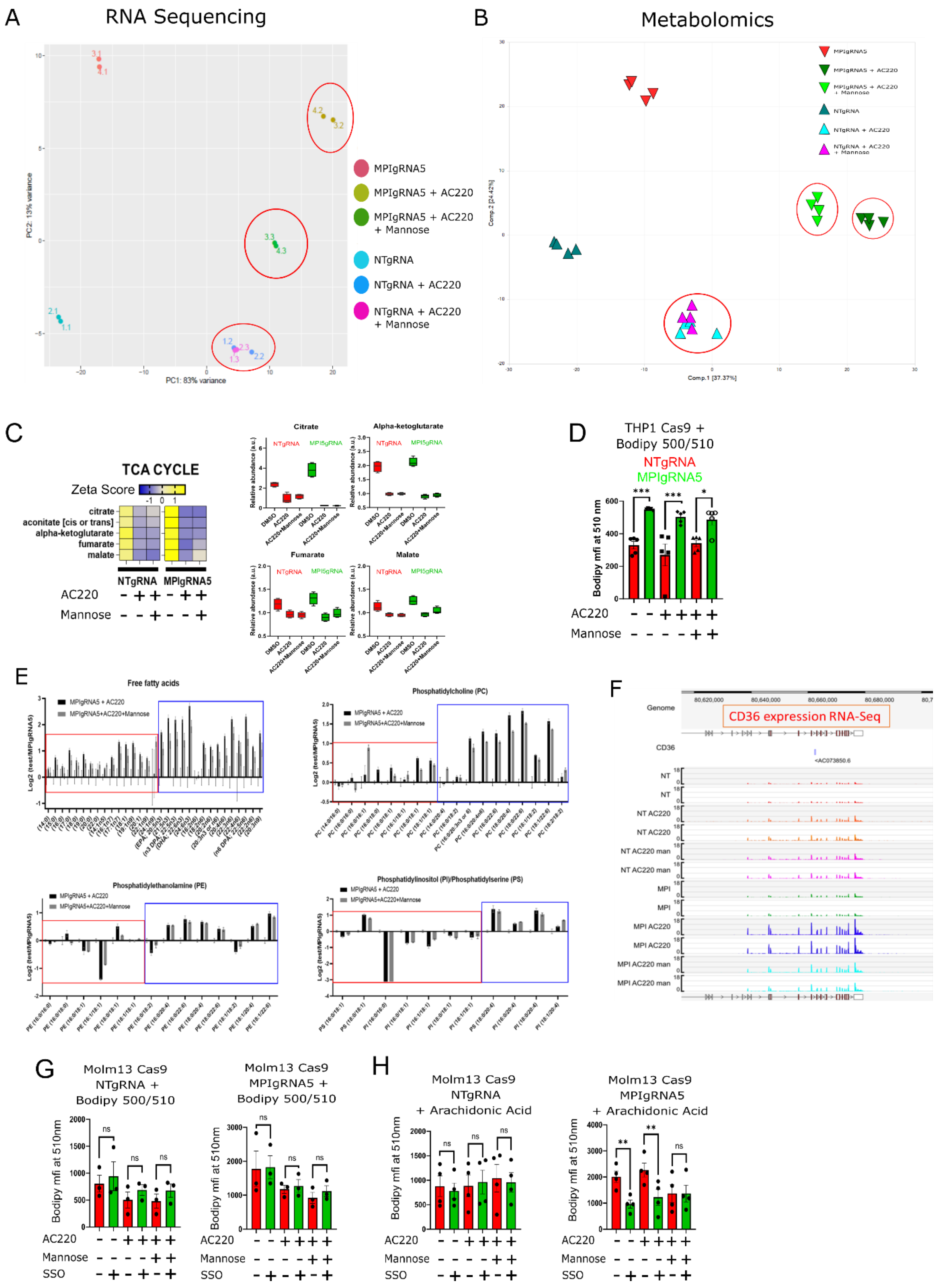
A
RNA Sequencing
PC1: 83% variance
PC2: 13% variance
MPIgRNA5
MPIgRNA5 + AC220
MPIgRNA5 + AC220 + Mannose
NTgRNA
NTgRNA + AC220
NTgRNA + AC220 + Mannose
B
Metabolomics
C
TCA CYCLE
Zeta Score
citrate
aconitate [cis or trans]
alpha-ketoglutarate
fumarate
malate
NTgRNA
MPIgRNA5
AC220
Mannose
Citrate
Alpha-ketoglutarate
Fumarate
Malate
D
THP1 Cas9 +
Bodipy 500/510
NTgRNA
MPIgRNA5
Bodipy mfi at 510 nm
E
Free fatty acids
Phosphatidylcholine (PC)
Phosphatidylethanolamine (PE)
Phosphatidylinositol (PI)/Phosphatidylserine (PS)
F
CD36 expression RNA-Seq
G
Molm13 Cas9
NTgRNA +
Bodipy 500/510
Molm13 Cas9
MPIgRNA5 +
Bodipy 500/510
H
Molm13 Cas9
NTgRNA
+ Arachidonic Acid
Molm13 Cas9
MPIgRNA5
+ Arachidonic Acid
SSO

**Extended data figure 3**
**Increased FA uptake in MPI KO is not exclusively caused by CD36**

**A** – Principal component analysis of RNA sequencing data from NT gRNA and MPI gRNA5 MOLM13 cells treated with vehicle, AC220 (1nM), mannose (100µM) or in combinations as indicated for 24 hours. Red circles indicate NT gRNA with AC220 and AC220 and mannose, MPI gRNA5 with AC220 and AC220 and mannose; **B** - Principal component analysis of global metabolomics data from NT gRNA and MPI gRNA5 MOLM13 cells treated with vehicle, AC220 (1nM) or mannose (100µM) as indicated for 48 hours. Red circles indicate NT gRNA with AC220 and AC220 and mannose, MPI gRNA5 with AC220 and AC220 and mannose; **C** – Comparison of levels of TCA cycle metabolites from global metabolic profiling of MOLM13 NT gRNA and MPI gRNA5 cells treated with vehicle, AC220 (1nM),mannose (100µM) or in combinations as indicated for 24 hours with heat map (left) and graphs of individual metabolites (right); **D** - Uptake of the fluorescent C1-Bodipy 500/510 C12 by NT gRNA or MPI gRNA5 THP1 cells treated with vehicle, AC220 (1nM) or AC220 (1nM) and mannose (100µM) for 24 hours, with a representative flow cytometry plot (right). N=5, 1 way Anova with Tukey's correction for multiple comparisons; **E** – Levels of free fatty acid (top left), phosphatidylcholine (top right) phosphatidylethanolamine (bottom left) and phosphatidylinositol/phosphatidylserine (bottom right) from global metabolic profiling in MPI gRNA5 cells treated with AC220 (1nM) or AC220 (1nM) and mannose (100µM) for 48 hours. Saturated or monounsaturated fatty acids are in the red boxes, while polyunsaturated fatty acids are in the blue boxes; **F** – Expression of CD36 RNA from RNA sequencing in NT gRNA and MPI gRNA5 MOLM13 cells treated with vehicle, AC220 (1nM), mannose (100µM) or in combinations as indicated for 48 hours. ; **G** - Uptake of the fluorescent C1-Bodipy 500/510 C12 by NT gRNA (left panel) or MPI gRNA5 (right panel) MOLM13 cells treated with SSO (100µM) or vehicle for 24 hours then vehicle, AC220 (1nM), mannose (100µM) or in combinations as indicated with C1-Bodipy 500/510 C12 for a further 24 hours. N=3, 1 way Anova with Tukey's correction for multiple comparisons; **H** - Uptake of arachidonic acid by NT gRNA (left panel) or MPI gRNA5 (right panel) MOLM13 cells treated with SSO (100µM) or vehicle for 24 hours then vehicle, AC220 (1nM), mannose (100µM) or in combinations as indicated with arachidonic acid for a further 24 hours. N=4, 1 way Anova with Tukey's correction for multiple comparisons. For all panels, ns = not significant, *=p<0.05, **=p<0.01, ***=p<0.005, ****=p<0.001.

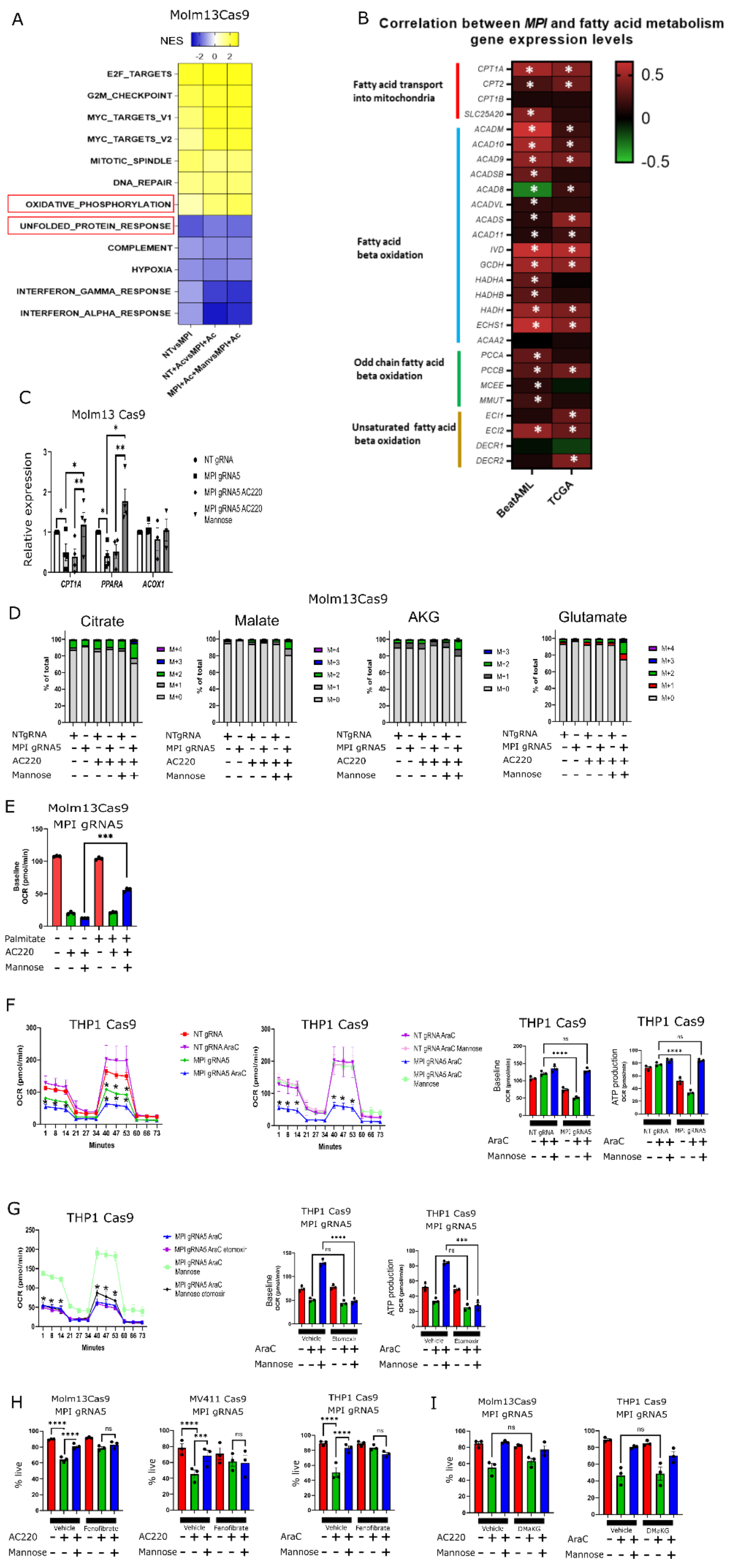
A
Molm13Cas9
NES
E2F_TARGETS
G2M_CHECKPOINT
MYC_TARGETS_V1
MYC_TARGETS_V2
MITOTIC_SPINDLE
DNA_REPAIR
OXIDATIVE_PHOSPHORYLATION
UNFOLDED_PROTEIN_RESPONSE
COMPLEMENT
HYPOXIA
INTERFERON_GAMMA_RESPONSE
INTERFERON_ALPHA_RESPONSE
NTvsMPI
NT+AcvsMPI+Ac
MPI+Ac+ManvsMPI+Ac
B
Correlation between MPI and fatty acid metabolism gene expression levels
Fatty acid transport into mitochondria
Fatty acid beta oxidation
Odd chain fatty acid beta oxidation
Unsaturated fatty acid beta oxidation
BeatAML
TCGA
C
Molm13 Cas9
Relative expression
NT gRNA
MPI gRNA5
MPI gRNA5 AC220
MPI gRNA5 AC220 Mannose
CPT1A
PPARA
ACOX1
Molm13Cas9
D
Citrate
Malate
AKG
Glutamate
% of total
NTgRNA
MPI gRNA5
AC220
Mannose
E
Molm13Cas9
MPI gRNA5
Baseline OCR (pmol/min)
Palmitate
AC220
Mannose
F
THP1 Cas9
OCR (pmol/min)
Minutes
NT gRNA
NT gRNA AraC
MPI gRNA5
MPI gRNA5 AraC
NT gRNA AraC Mannose
MPI gRNA5 AraC Mannose
Baseline OCR (pmol/min)
ATP production OCR (pmol/min)
AraC
Mannose
G
THP1 Cas9
MPI gRNA5 AraC etomoxir
MPI gRNA5 AraC Mannose
MPI gRNA5 AraC Mannose etomoxir
THP1 Cas9
MPI gRNA5
Vehicle
Etomoxir
AraC
Mannose
H
Molm13Cas9
MPI gRNA5
MV411 Cas9
MPI gRNA5
THP1 Cas9
MPI gRNA5
% live
Vehicle
Fenofibrate
AC220
AraC
Mannose
I
Molm13Cas9
MPI gRNA5
THP1 Cas9
MPI gRNA5
DMαKG

**Extended data figure 4**
**MPI KO causes a reduction in fatty acid oxidation in AML cell lines**

**A** – Heatmap showing NES for the most significantly up or downregulated hallmark genesets from RNA sequencing comparing NT gRNA or MPI gRNA5 treated with AC220 (1nM), mannose (100µM) or in combinations (highlighted in red pathways relevant to the mechanism described in the manuscript); **B** – Heatmap showing correlation between *MPI* and fatty acid metabolism genes expression from the BeatAML (left column) and TCGA (right column) AML databases. *=significant correlation, Spearman's Rank correlation; **C** – Relative expression of *CPT1A*, *PPARA* and *ACOX1*, normalised to *ACTB* expression, from RT-qPCR, in NT gRNA MOLM13 cells treated with Vehicle and MPI gRNA5 MOLM13 cells treated with vehicle, AC220 (1nM) or AC220 (1nM) and mannose (100µM) for 72 hours. *ACOX1* was chosen as a control gene involved with peroxisomal lipid oxidation. N=4, 1 way Anova with Tukey's correction for multiple comparisons. **D** - Percentage of TCA cycle intermediates and associated metabolites labelled with $^{13}C$ from $^{13}C_{16}$-palmitate, Citrate (left), Malate ($2^{nd}$ left), Alpha-ketoglutarate ($2^{nd}$ right) and Glutamate (far right) in MPI KO and NT MOLM13 cells treated with AC220 (1nm) and mannose (100µM) as indicated for 24 hours along with 50µM $^{13}C$-palmitate. Includes all labelled and unlabelled species detected. N=5; **E** - Baseline OCR of MPI gRNA5 cells cultured overnight in substrate limited RPMI media without FBS and glutamine treated with palmitate (50µM), vehicle, AC220 (1nM), mannose (100µM) in combinations as indicated from SEAHORSE MitoStress test. N=3, 1 way Anova with Tukey's correction for multiple comparisons; **F**- SEAHORSE MitoStress tests showing oxygen consumption rate over time comparing NT gRNA and MPI gRNA5 THP1 cells treated with vehicle, AraC (1µM) or AraC (1µM) and mannose (100µM) as indicated after 72 hours of treatment, N=3, 2 way Anova with Sidak's correction for multiple comparisons (two left panels). Baseline OCR and ATP production of NT gRNA and MPI gRNA5 THP1 cells treated with vehicle, AraC (1µM)) or AraC (1µM) and mannose (100µM) as indicated after 72 hours of treatment, N=3, 1 way Anova with Tukey's correction for multiple comparisons (2 right panels); **G** - SEAHORSE MitoStress tests showing oxygen consumption rate over time comparing MPI gRNA5 THP1 cells treated with AraC (1µM), mannose (100µM) after 72 hours in combinations as indicated with or without etomoxir (50µM), N=3, 2 way Anova with Sidak's correction for multiple comparisons (left panel). Baseline OCR and ATP production of MPI gRNA5 THP1 cells treated with vehicle, AraC (1µM), mannose (100µM) after 72 hours in combinations as indicated with or without etomoxir (50µM), N=3, 1 way Anova with Tukey's correction for multiple comparisons (2 right panels); **H** - Percentage of live MPI gRNA5 MOLM13 cells (left), MV411 (middle) and THP1 (right) treated with vehicle, fenofibrate (10µM), AC220 (1nM), mannose (100µM) or in combinations as indicated 72 hours after treatment. N=3, 1 way Anova with Tukey's correction for multiple comparisons; **I** – Percentage of live MPI gRNA5 MOLM13 cells (left) and THP1 (right) treated with vehicle, Dimethyl Alpha-ketoglutarate (DMaKG, 4mM), AC220 (1nM), mannose (100µM) or in combinations as indicated 72 hours after treatment. N=3, 1 way Anova with Tukey's correction for multiple comparisons. For all panels, ns = not significant, *=p<0.05, **=p<0.01, ***=p<0.005, ****=p<0.001.

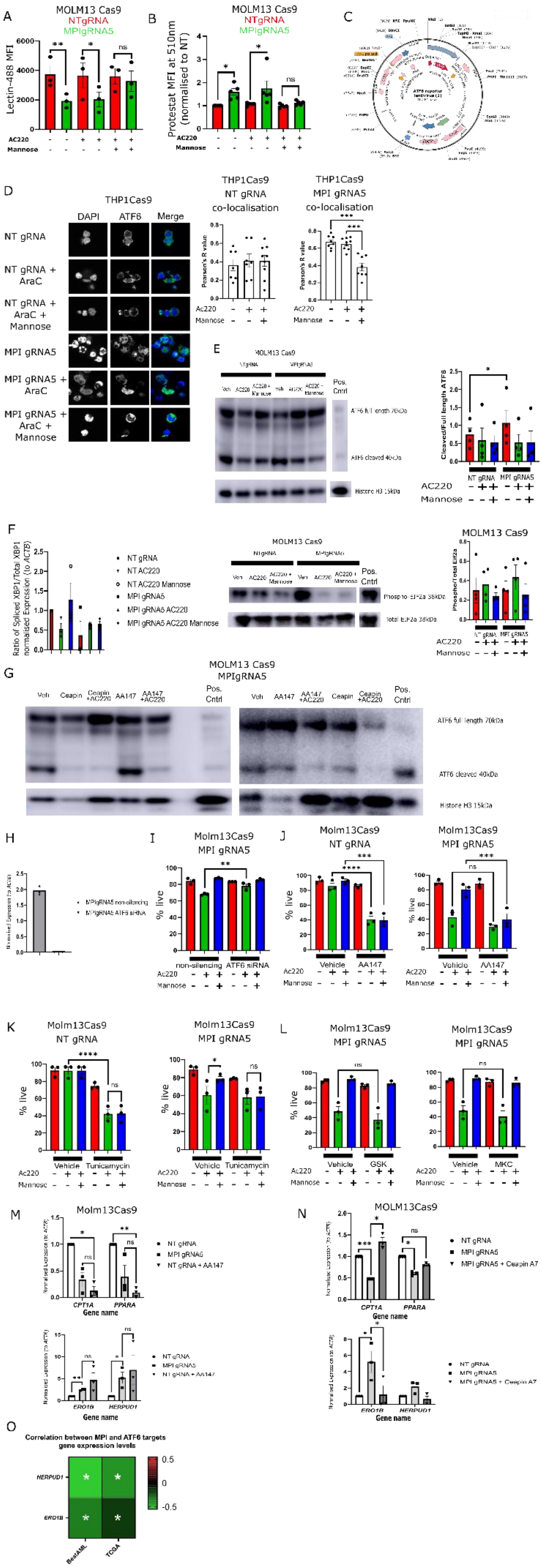
A
MOLM13 Cas9
NTgRNA
MPIgRNA5
Lectin-488 MFI
AC220
Mannose
B
Protestat MFI at 510nm (normalised to NT)
C
D
THP1Cas9
DAPI
ATF6
Merge
NT gRNA
NT gRNA + AraC
NT gRNA + AraC + Mannose
MPI gRNA5
MPI gRNA5 + AraC
MPI gRNA5 + AraC + Mannose
THP1Cas9 NT gRNA co-localisation
THP1Cas9 MPI gRNA5 co-localisation
Pearson's R value
E
Pos. Cntrl
Histone H3 15kDa
Cleaved/Full length ATF6
NT gRNA
MPI gRNA5
F
NT gRNA
NT AC220
NT AC220 Mannose
MPI gRNA5
MPI gRNA5 AC220
MPI gRNA5 AC220 Mannose
Total EIF2a 38kDa
Phospho/Total Eif2a
G
MOLM13 Cas9 MPIgRNA5
Veh
Ceapin
Ceapin +AC220
AA147
AA147 +AC220
ATF6 full length 70kDa
ATF6 cleaved 40kDa
Histone H3 15kDa
H
I
Molm13Cas9 MPI gRNA5
% live
non-silencing
ATF6 siRNA
J
Molm13Cas9 NT gRNA
Vehicle
AA147
K
Tunicamycin
L
GSK
MKC
M
Molm13Cas9
NT gRNA
MPI gRNA5
NT gRNA + AA147
CPT1A
PPARA
ERO1B
HERPUD1
Gene name
N
MOLM13Cas9
MPI gRNA5 + Ceapin A7
O
Correlation between MPI and ATF6 targets gene expression levels
HERPUD1
ERO1B
BeatAML
TCGA

**Extended data figure 5**
**ATF6 arm of the UPR is increased in MPI KO AML cells and contributes to cell death**

**A** – Lectin-488 MFI from flow cytometry of NT gRNA or MPI gRNA5 MOLM13 cells treated with vehicle, AC220 (1nM) or AC220 (1nM) and Mannose (100µM) for 72 hours. N=3, 1 way Anova with Tukey's correction for multiple comparisons; **B –** Proteostat MFI from flow cytometry of NT gRNA or MPI gRNA5 MOLM13 cells treated with vehicle, AC220 (1nM) or AC220 (1nM) and Mannose (100µM) for 72 hours. N=3, 1 way Anova with Tukey's correction for multiple comparisons; **C** – Plasmid map of the YFP-linked ATF6 reporter; **D** – Confocal microscopy images of NT gRNA and MPI gRNA5 THP1 cells stained for DAPI (blue, 1st column) and ATF6 (green, 2nd column) with a merged image (3rd column), treated with vehicle, AraC (1µM), mannose (100µM) or in combinations as indicated 72 hours after treatment (left). Colocalisation analysis from immunofluorescence images of NTgRNA (left) and MPIgRNA5 (right) THP1 cells treated with vehicle, AraC (1µM), mannose (100µM) or in combinations as indicated. Analysis performed with Coloc2 plugin in ImageJ, ordinary 1 way Anova with Tukey's correction for multiple comparisons (right); **E** - Western blot of protein samples from NT gRNA or MPI gRNA5 MOLM13 cells treated with vehicle, AC220 (1nM) or AC220 (1nM) and mannose (100µM) as indicated at 72 hours of treatment. Primary antibodies ATF6 and total Histone H3 as a loading control (Left). Densitometry of cleaved ATF6 divided full length ATF6 (right); **F -** Relative expression of spliced *XBP1* and total *XBP1* normalised to *ACTB* expression, from RT-qPCR, in NT gRNA and MPI gRNA5 MOLM13 cells treated with vehicle, AC220 (1nM) or mannose (100µM) in combinations as indicated for 72 hours (left panel). Western blot of protein samples from NT gRNA or MPI gRNA5 MOLM13 cells treated with vehicle, AC220 (1nM) or AC220 (1nM) and mannose (100µM) as indicated at 72 hours of treatment. Primary antibodies EIF2α and phospho-EIF2α (middle panel) with densitometry analysis of phospho EIF2α divided by total EIF2α (right panel). N=3; **G –** Western blot of protein samples from NT gRNA or MPI gRNA5 MOLM13 cells treated with vehicle, AC220 (1nM), AA147 (5µM), Ceapin A7 (1µM) or in combinations as indicated at 72 hours of treatment. Primary antibodies ATF6 and total Histone H3 as a loading control; **H –** Relative expression of *ATF6* to *ACTB* expression, from RT-qPCR, in MPI gRNA5 MOLM13 cells transduced with 10nM ATF6 or non-silencing control siRNA; **I** - Percentage of live cells of MPI gRNA5 MOLM13 cells transfected with 10nM ATF6 or non-silencing siRNA for 48 hours then treated with vehicle, AC220 (1nM) or AC220 (1nM) and mannose (100µM) as indicated for 72 hours (right panel). One way Anova with Tukey's correction for multiple comparisons. **J** - Percentage of live cells of NT gRNA (left) or MPI gRNA5 (right) MOLM13 cells treated with vehicle, AC220 (1nM), mannose (100µM), AA147 (5µM) or in combinations as indicated 72 hours after treatment. N=3, 1 way Anova with Tukey's correction for multiple comparisons; **K** - Percentage of live cells of NT gRNA (left) and MPI gRNA5 (right) MOLM13 cells treated with vehicle, AC220 (1nM), mannose (100µM), tunicamycin (1µM) or in combinations as indicated 72 hours after treatment (left). N=3, 1 way Anova with Tukey's correction for multiple comparisons; **L -** Percentage of live cells of MPI gRNA5 MOLM13 cells treated with vehicle, AC220 (1nM), mannose (100µM), GSK2656157 (200nM, left panel), MKC-3946 (1µM, right panel) or in combinations as indicated 72 hours after treatment . N=3, ns=not significant, 1 way Anova with Tukey's correction for multiple comparisons; **M** - Relative expression of *CPT1A* and *PPARA* (top) and *ERO1B* and *HERPUD1* (bottom) normalised to *ACTB* expression, from RT-qPCR, in NT gRNA and MPI gRNA5 MOLM13 cells treated with vehicle or AA147 (5µM) for 72 hours. N=3, 1 way Anova with Tukey's correction for multiple comparisons; **N** - Relative expression of *CPT1A* and *PPARA* (top) and *ERO1B* and *HERPUD1* (bottom) normalised to *ACTB* expression, from RT-qPCR, in NT gRNA and MPI gRNA5 MOLM13 cells treated with vehicle or CeapinA7 (1µM) for 72 hours. N=3, 1 way Anova with Tukey's correction for multiple comparisons; **O** – Heatmap showing correlation between *MPI* and ATF6 target genes (*ERO1B* and *HERPUD1*) expression from the BeatAML (left column) and TCGA (right column) AML databases. Spearman's Rank correlation. For all panels, ns = not significant, *=p<0.05, **=p<0.01, ***=p<0.005, ****=p<0.001.

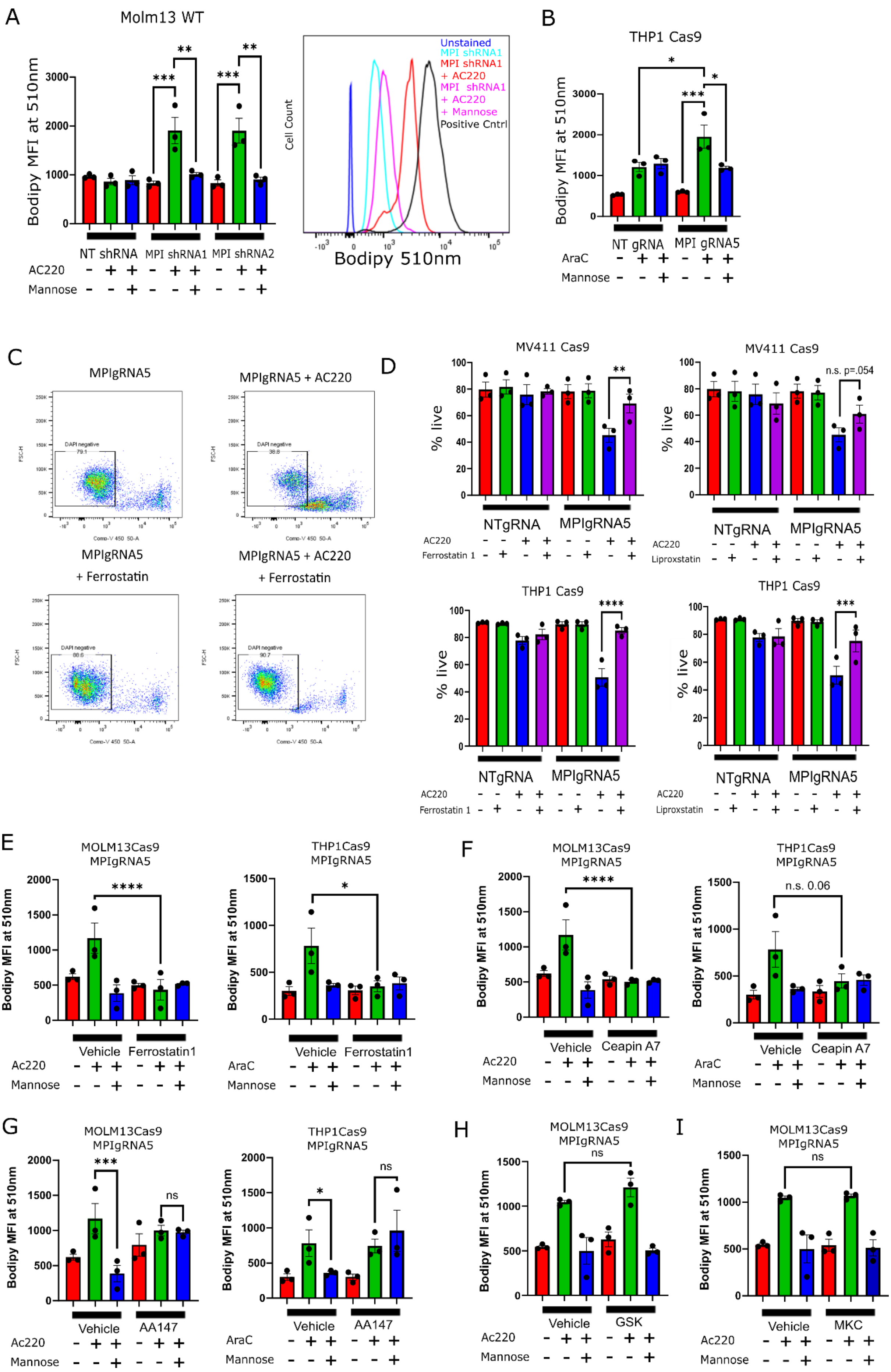

A
Molm13 WT
Bodipy MFI at 510nm
NT shRNA
MPI shRNA1
MPI shRNA2
AC220
Mannose
Cell Count
Unstained
MPI shRNA1
MPI shRNA1 + AC220
MPI shRNA1 + AC220 + Mannose
Positive Cntrl
Bodipy 510nm
B
THP1 Cas9
NT gRNA
MPI gRNA5
AraC
C
MPIgRNA5
MPIgRNA5 + AC220
MPIgRNA5 + Ferrostatin
MPIgRNA5 + AC220 + Ferrostatin
DAPI negative
D
MV411 Cas9
THP1 Cas9
% live
NTgRNA
MPIgRNA5
Ferrostatin 1
Liproxstatin
n.s. p=.054
E
MOLM13Cas9 MPIgRNA5
THP1Cas9 MPIgRNA5
Vehicle
Ferrostatin1
Ac220
F
Ceapin A7
n.s. 0.06
G
AA147
ns
H
GSK
I
MKC

**Extended data figure 6**
**MPI KO AML cell lines are primed for ferroptosis via sensitivity to PUFAs**

**A** - Mean fluorescence intensity at 510nm of Bodipy 581/591 C11, which shows level of lipid peroxidation, in NT shRNA, MPI shRNA1 and MPI shRNA2 MOLM13 cells treated with vehicle, AC220 (1nM) or mannose (100µM) in the indicated combinations after 72 hours of treatment (left) with a representative flow cytometry plot (right including positive control). N=3, 1 way Anova with Tukey's correction for multiple comparisons; **B** - Mean fluorescence intensity at 510nm of Bodipy 581/591 C11, which shows level of lipid peroxidation, in NTgRNA and MPIgRNA5 THP1 cells treated with AraC (1µM), mannose (100µM) or in combinations as indicated 72 hours after treatment. N=3, 1 way Anova with Tukey's correction for multiple comparisons; **C** – Representative flow cytometry plots showing DAPI intensity vs FSC-H from MPI gRNA5 MOLM13 cells treated with AC220 (1nM) and ferrostatin-1 (5µM) in combinations as indicated for 6 days; **D -** Percentage of live NT gRNA and MPI gRNA5 MV411 (top panels) or THP1 (bottom panels) cells treated with vehicle, AC220 (1nM), AraC (1µM), mannose (100µM), ferrostatin-1 (5µM, left), liproxstatin (1µM, right) or combinations as indicated for 6 days. N=3, 1 way Anova with Tukey's correction for multiple comparisons. **E -** Mean fluorescence intensity at 510nm of Bodipy 581/591 C11, which shows level of lipid peroxidation, in MPI gRNA5 MOLM13 (left) and THP1 (right) cells treated with AC220 (1nM), AraC (1µM), mannose (100µM), ferrostatin1 (5µM) or in combinations as indicated 72 hours after treatment. N=3, 1 way Anova with Tukey's correction for multiple comparisons; **F -** Mean fluorescence intensity at 510nm of Bodipy 581/591 C11, which shows level of lipid peroxidation, in MPI gRNA5 MOLM13 (left) and THP1 (right) cells treated with AC220 (1nM), AraC (1µM), mannose (100µM), Ceapin A7 (1µM) or in combinations as indicated 72 hours after treatment. N=3, 1 way Anova with Tukey's correction for multiple comparisons; **G -** Mean fluorescence intensity at 510nm of Bodipy 581/591 C11, which shows level of lipid peroxidation, in MPI gRNA5 MOLM13 (left) and THP1 (right) cells treated with AC220 (1nM), AraC (1µM), mannose (100µM), AA147 (5µM) or in combinations as indicated 72 hours after treatment. N=3, 1 way Anova with Tukey's correction for multiple comparisons; **H -** Mean fluorescence intensity at 510nm of Bodipy 581/591 C11, which shows level of lipid peroxidation, in MPI gRNA5 MOLM13 cells treated with AC220 (1nM), mannose (100µM), GSK2656157 (200nM) or in combinations as indicated 72 hours after treatment. N=3, 1 way Anova with Tukey's correction for multiple comparisons; **I -** Mean fluorescence intensity at 510nm of Bodipy 581/591 C11, which shows level of lipid peroxidation, in MPI gRNA5 MOLM13 cells treated with AC220 (1nM), mannose (100µM), MKC-3946 (1µM) or in combinations as indicated 72 hours after treatment. N=3, 1 way Anova with Tukey's correction for multiple comparisons. For all panels, ns = not significant, *=p<0.05, **=p<0.01, ***=p<0.005, ****=p<0.001.

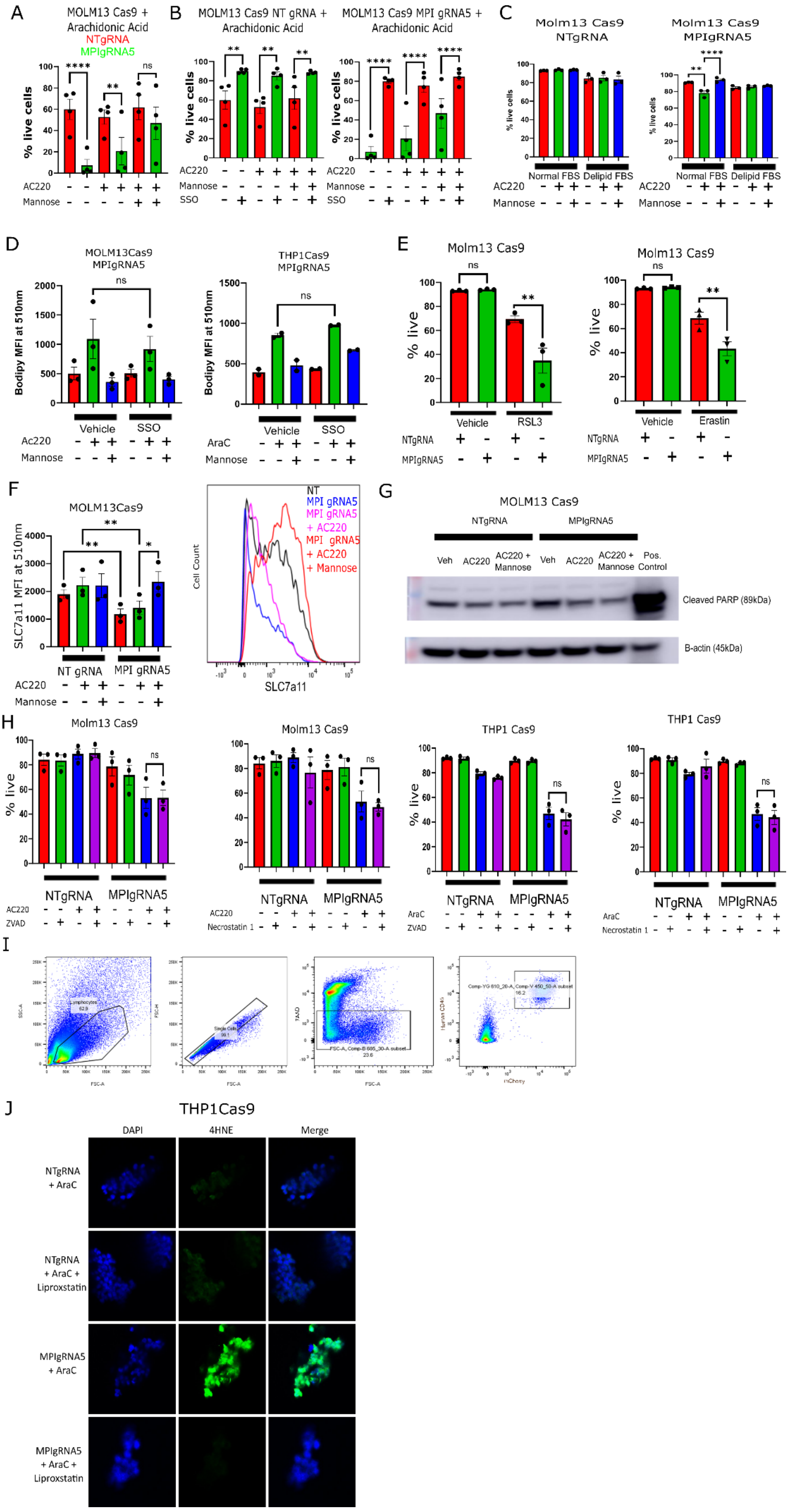
A
MOLM13 Cas9 + Arachidonic Acid
NTgRNA
MPIgRNA5
% live cells
AC220
Mannose
B
MOLM13 Cas9 NT gRNA + Arachidonic Acid
MOLM13 Cas9 MPI gRNA5 + Arachidonic Acid
% live cells
AC220
Mannose
SSO
C
Molm13 Cas9 NTgRNA
Molm13 Cas9 MPIgRNA5
% live cells
Normal FBS
Delipid FBS
AC220
Mannose
D
MOLM13Cas9 MPIgRNA5
THP1Cas9 MPIgRNA5
Bodipy MFI at 510nm
Vehicle
SSO
Ac220
AraC
Mannose
E
Molm13 Cas9
% live
Vehicle
RSL3
Erastin
NTgRNA
MPIgRNA5
F
MOLM13Cas9
SLC7a11 MFI at 510nm
NT gRNA
MPI gRNA5
AC220
Mannose
NT
MPI gRNA5
MPI gRNA5 + AC220
MPI gRNA5 + AC220 + Mannose
Cell Count
SLC7a11
G
MOLM13 Cas9
NTgRNA
MPIgRNA5
Veh
AC220
AC220 + Mannose
Pos. Control
Cleaved PARP (89kDa)
B-actin (45kDa)
H
Molm13 Cas9
THP1 Cas9
% live
NTgRNA
MPIgRNA5
AC220
AraC
ZVAD
Necrostatin 1
I
J
THP1Cas9
DAPI
4HNE
Merge
NTgRNA + AraC
NTgRNA + AraC + Liproxstatin
MPIgRNA5 + AraC
MPIgRNA5 + AraC + Liproxstatin

**Extended data figure 7**
**MPI KO AML cell lines are sensitive to ferroptotic cell death as a result of ATF6 activation**

**A** - Percentage of NT gRNA and MPI gRNA5 MOLM13 live cells treated with arachidonic acid (1µM) in combination with vehicle, AC220 (1nM), mannose (100µM) or combinations as indicated for 48 hours. N=4, ns = not significant, 1 way Anova with Tukey's correction for multiple comparisons; **B** - Percentage of NT gRNA (left graph) and MPI gRNA5 (right graph) MOLM13 live cells when treated with SSO (100µM) or vehicle for 24 hours then arachidonic acid (1µM) in combination with vehicle, AC220 (1nM), mannose (100µM) or in combinations as indicated for a further 48 hours; N=4, 1 way Anova with Tukey's correction for multiple comparisons **C** - Percentage of live cells of NT gRNA (left) or MPI gRNA5 (right) MOLM13 cells cultured in either normal FBS or delipidated FBS and treated with vehicle, AC220 (1nM), mannose (100µM) or in combinations as indicated 72 hours after treatment. N=3, ns=not significant, 1 way Anova with Tukey's correction for multiple comparisons; **D** - Mean fluorescence intensity at 510nm of Bodipy 581/591 C11, which shows level of lipid peroxidation, in MPI gRNA5 MOLM13 (left) and THP1 (right) cells treated with AC220 (1nM), AraC (1µM), mannose (100µM), SSO (100µM) or in combinations as indicated 72 hours after treatment. N=3, 1 way Anova with Tukey's correction for multiple comparisons; **E –** Percentage of live NT gRNA and MPI gRNA5 Molm13 cells treated with RSL3 (100nM,left panel) or erastin (4µM, right panel) after 48 hours. N=3, ns = not significant, 1 way Anova with Tukey's correction for multiple comparisons; **F -** Mean fluorescence intensity from flow cytometry analysis of SLC7a11 on the surface of MOLM13 NT gRNA and MPI gRNA5 cells treated with vehicle, AC220 (1nM), mannose (100µM) or in combinations as indicated after 24 hours of treatment, N=3, *=p<0.05, **=p<0.01, 1 way Anova with Tukey's correction for multiple comparisons (left). Representative plots of flow cytometry fluorescence intensity plot of MOLM13 NT gRNA and MPI gRNA5 cells with vehicle, AC220 (1nM), mannose (100µM) or in combinations as indicated (right); **G -** Representative western blot of cleaved PARP in MOLM13 NTgRNA and MPIgRNA5 treated with AC220 (1nM) and mannose (100µM) in combinations as indicated for 6 days. **H -** Percentage of NT gRNA and MPI gRNA5 MOLM13 (2 left panels) or THP1 (2 right panels) live cells treated with vehicle, AC220 (1nM), mannose (100µM), ZVAD (20µM), necrostatin (20µM) or combinations as indicated for 72 hours. N=3, 1 way Anova with Tukey's correction for multiple comparisons; **I -** Representative flow cytometry plots of the sorting of THP1 cells from transplanted NBSGW mice; **J** – Representative immunofluorescence images from NT gRNA or MPI gRNA5 THP1 cells sorted from transplanted NBSGW mice treated with AraC (50mg/kg, 7 days), liproxstatin (10mg/kg, 10 days), or in combinations as indicated. DAPI (left column), 4HNE (middle column) and a merged image (right column). For all panels, ns = not significant, *=p<0.05, **=p<0.01, ***=p<0.005, ****=p<0.001.

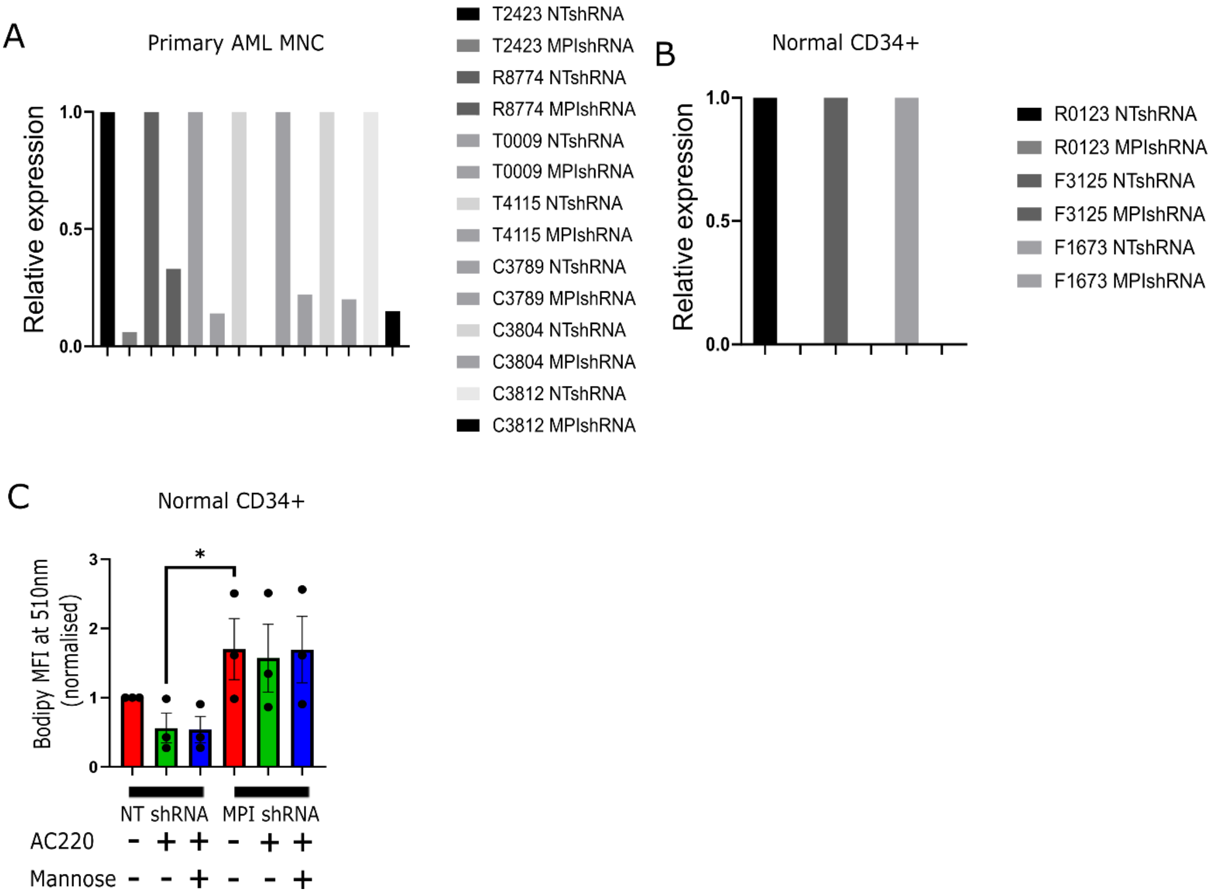


**Extended data figure 8**

**MPI knockdown efficiency in primary AML and CD34 samples**

**A** – Relative *MPI* expression measured by RT-qPCR in primary AML $FLT3^{ITD}$ samples transduced with either NT shRNA or MPI shRNA1; **B** - Relative *MPI* expression measured by RT-qPCR in normal CD34+ samples transduced with either NT shRNA or MPI shRNA1; **C** - Mean fluorescence intensity at 510nm of Bodipy 581/591 C11, which shows level of lipid peroxidation, in NT shRNA and MPI shRNA1 normal CD34+ cells treated with vehicle, AC220 (2.5nM) and mannose (100µM) in the indicated combinations at 72 hours after treatment. N=3 for each sample, 1 way Anova with Tukey's correction for multiple comparisons. For all panels, ns = not significant, *=p<0.05, **=p<0.01, ***=p<0.005, ****=p<0.001.
